# Spatial and temporal regulation of Wnt signaling pathway members in the development of butterfly eyespots

**DOI:** 10.1101/2023.04.13.536826

**Authors:** Tirtha Das Banerjee, Suriya Narayanan Murugesan, Antόnia Monteiro

## Abstract

Wnt signaling is involved in the differentiation of eyespot color patterns on the wings of butterflies, but the identity and spatio-temporal regulation of specific Wnt pathway members remains unclear. Here we explore the localization and function of armadillo/β-catenin dependent (canonical) and armadillo/β- catenin independent (non-canonical) Wnt signaling in eyespot development in *Bicyclus anynana* by localizing Armadillo (Arm), the expression of all seven *wnt* ligand and four *frizzled* receptor transcripts present in the genome of this species, and testing the function of *arm* and *frizzled4* using CRISPR-Cas9. During mid to late larval wing development, Arm protein was present in cells at the center of the future eyespots, the foci, and the wing margin, but *wnts* expressed on the wing, *wnt1* (*wingless*), *wnt6*, and *wnt10* showed expression only some distance away from the foci, along the wing margin. The receptor *frizzled9* was expressed in the wing margin and in finger-like projections leading to the foci during early larval wing development, overlapping in expression with Arm. At the same time, the receptor *frizzled4* showed a novel expression pattern, anti-localized with Arm, where it is likely transducing non-canonical Wnt signaling. In the early pupal stage, *wnt1* was newly expressed in the foci, as previously shown, along with Arm. In addition, *frizzled4* and *frizzled9*-mediated Wnt signaling is likely repressing the expression of *frizzled2*, as these receptors have anti-colocalized expression domains. Arm had a conserved expression in three other nymphalid butterflies, and functional knockouts of *arm* and *frizzled4* in *B. anynana* showed that both genes are essential for the differentiation of eyespots. These results show that distinct Wnt signaling pathways are essential for eyespot development in butterflies and are likely interacting to control their active domains.

## Introduction

Wnt signaling is fundamental to cellular communication in multicellular organisms. This communication involves secreted glycoproteins, the Wnt ligands, that are produced in signaling cells, traveling some distance away to regulate the expression of target genes in surrounding cells (Neumann & Cohen, 1997; Wiese et al., 2018). Understanding the mechanisms underlying Wnt signaling is fundamental to studies of both normal and altered development, such as in wound healing and cancer, and has remained elemental in basic and applied biological research (Lawrence et al., 1996; P. Liu et al., 1999; McMahon & Bradley, 1990; Van Camp et al., 2014; Polakis, 2007; Whyte et al., 2012). There is, however, considerable debate about the spatial and temporal regulation and interaction of different Wnt signaling pathways; the distance and the mechanisms by which Wnts can travel across tissues; the mechanism by which distinct Wnt pathway members are regulated in cells; and how different Wnt ligands and receptors interact with each other (Cox & Peifer, 1998; Routledge & Scholpp, 2019; Van Amerongen et al., 2008).

In classical studies there are two main pathways involved in Wnt signaling – a canonical and a non-canonical pathway - which use different ligand and receptor gene paralogues to transduce extra-cellular signals. In canonical Wnt signaling in *Drosophila* (**Figure S1A, B**) specific Wnt ligands bind to specific Frizzled receptors at the cell surface, which then signal through Armadillo (β-catenin in vertebrates) to regulate gene expression in the nucleus (Bejsovec, 2006, 2013; Gao & Chen, 2010; J. Liu et al., 2022). Non-canonical Wnt signaling works independent of Arm/β-catenin and includes the planar cell polarity (PCP) pathway, which regulates the cytoskeleton of cells (**Figure S1C**), amongst other pathways (Ewen-Campen et al., 2020; Guo et al., 2004; Korswagen, 2002; Widelitz, 2005). PCP signaling has been reported to work both in a Wnt ligand-dependent and in a Wnt ligand-independent manner in mouse and *Drosophila,* respectively (Ewen-Campen et al., 2020; Mattes et al., 2018; Yu et al., 2020). Both canonical and non-canonical pathways, however, are involved in similar processes of tissue organization, cell proliferation, and cell-cell communication (Butler & Wallingford, 2017; Goodrich & Strutt, 2011; Nusse, 2006; Van Amerongen et al., 2008; Wiese et al., 2018).

The distinct Wnt ligands and Frizzled receptors used by both pathways belong to multi-copy gene families. In insect genomes there are six to nine *wnts* (Mundaca-Escobar et al., 2022) and three to four *frizzled* genes (Janssen et al., 2015), while in mammalian systems there are 19 *wnts* and 10 *frizzled* genes (Van Amerongen et al., 2008; Widelitz, 2005). Classically, *wnts* such as *wnt1*, *wnt3*, *wnt8*, and *wnt10* have been associated with the canonical pathway, while *wnt5a*, *wnt7a*, and *wnt11* with the non-canonical pathway in metazoans (Heisenberg et al., 2000; Le Grand et al., 2009; Tada & Smith, 2000; Van Amerongen et al., 2008; Wallingford et al., 2001). Newer studies, however, have contradicted such categorization since some Wnts can transduce alternate Wnt signaling when coupled with specific receptors (Van Amerongen et al., 2008). Wnt5a, for example, can work both in the canonical pathway, using receptors such as *frizzled4*, and non-canonical pathway using *frizzled8* receptors (Mikels & Nusse, 2006; Wallingford et al., 2001).

Wnt signaling, using *wnt1*, has previously been implicated in regulating butterfly eyespot ring size, but the involvement of Wnt signaling in differentiating eyespot centers, critical for ring differentiation, has not been investigated. Multiple butterfly species have stable *wnt1* expression along the wing margin thoughout larval development (Carroll et al., 1994; Monteiro et al., 2006; Martin & Reed, 2010, 2014), but the signal transducer of canonical Wnt signaling, Armadillo (Arm), displays a dynamic pattern of expression during the larval stages (Banerjee & Monteiro, 2020; Connahs et al., 2019). Arm starts to be expressed across the whole wing but later resolves into broad vein and marginal expression. This expression then progressively narrows on top of the veins, wing margin, and also along fingers parallel to and centered between wing veins. These fingers contain an enlarged cluster of Arm-expressing cells in the middle, the eyespot foci, that mark the future eyespot centers (Banerjee & Monteiro, 2020; Connahs et al., 2019). Later in development, during the early pupal stage, *wnt1* is newly expressed in these focal cells suggesting a role in signaling from these cells to pattern the rings of color around the focus (Monteiro et al., 2006; Özsu et al., 2017). Transgenic RNAi studies against *wnt1*, where *wnt1* was down-regulated at the end of the pre-pupal stage and beginning of the pupal stage, resulted in reduction of all the colored rings, indicating that *wnt1* regulates eyespot size (Özsu et al., 2017). It is, however, still unclear whether Arm or Wnt signaling is required for eyespot center differentiation in the larval stages. Furthermore, the identity of the Wnt ligand(s) leading to Armadillo nuclear localization in the eyespot center during larval wing development is currently unknown, and we also lack information on the expression and function of any of the *frizzled* receptors in butterfly wings.

In the present work we explored the spatial and temporal expression of different Wnt signaling pathway genes in eyespot formation of *Bicyclus anynana.* We first focused on the localization of all the *wnt* ligands, *frizzled* receptors, Arm signal transducer, and Wnt target genes *distal-less* (*dll*) and *vestigial* (*vg*) in larval and pupal wings, and then we targeted the function of two of these genes, *arm* and *frizzled4,* with CRISPR-Cas9.

## Results

### Phylogenetic analysis of *wnt* and *frizzled* genes of *Bicyclus anynana*

To discover all possible *wnt* and *frizzled* genes present in the *B. anynana* genome we searched for the corresponding gene annotations in NCBI (*Bicyclus anynana* (taxid:110368)) and also blasted the orthologous *wnt* and *frizzled* sequences from *Drosophila melanogaster* in flybase against the *B. anynana* genome nBa.0.1, v1.2, and v2.0 in lepbase (Challis et al., 2016; Nowell et al. 2017; Murugesan et al. 2022). We identified seven *wnts* (excluding *wntA*), and four *frizzled* genes in the genome. *wnt9* was only found in nBa.0.1 but not v1.2 and v2.0 of the *B. anynana* genome (Holzem et al., 2019). A phylogenetic analysis performed with these genes and orthologues from other insects, showed that the seven *wnt*s represent *wnt1, wnt5, wnt6, wnt7, wnt9, wnt10,* and *wnt11,* who cluster with members of the same gene family from other butterfly and insect species (**Figure S2 and S3**). Similarly, a separate phylogenetic analysis identified *frizzled, frizzled2, frizzled4*, and *frizzled9* as being the four *frizzled* genes in the *B. anynana* genome (**Figure S4 and S5**).

### Armadillo (Arm) is expressed in a dynamic pattern in larval and pupal wings

Previous gene expression, transcriptomic, and modeling studies have proposed a role for Arm in the differentiation of the eyespot foci (Connahs et al., 2019; Banerjee and Monteiro, 2020; Wee et al., 2022). To extend these findings we used immunostainings to document Arm’s spatial-temporal expression across both larval and pupal wings at multiple time points, and across a longer period of wing development than previously investigated. In early fifth instar larvae, Arm was homogeneously distributed across the wing (stage 0.25, **Figure 1A**). As the wing developed, the protein localized along the vein cells, wing margin, and at the eyespot foci (stages 0.50-2.00, **Figure 1A**). During the pupal stage, Arm protein was observed in the eyespot foci and along the wing margin from 15-24 hrs after pupation (AP) (**Figure 1B**). We lack data from the early hours after pupation (prior to 15 hrs AP) because wings are too fragile to handle before this stage. Arm protein was also observed in the foci in three other nymphalid butterflies during mid-late larval wing development along with the eyespot marker protein Spalt (**Figure S6**). We conclude that Arm protein, and canonical Wnt signaling, is likely continuously present in the eyespot foci from mid fifth instar larval development to at least 24 hours after pupation.

**Figure 1.**
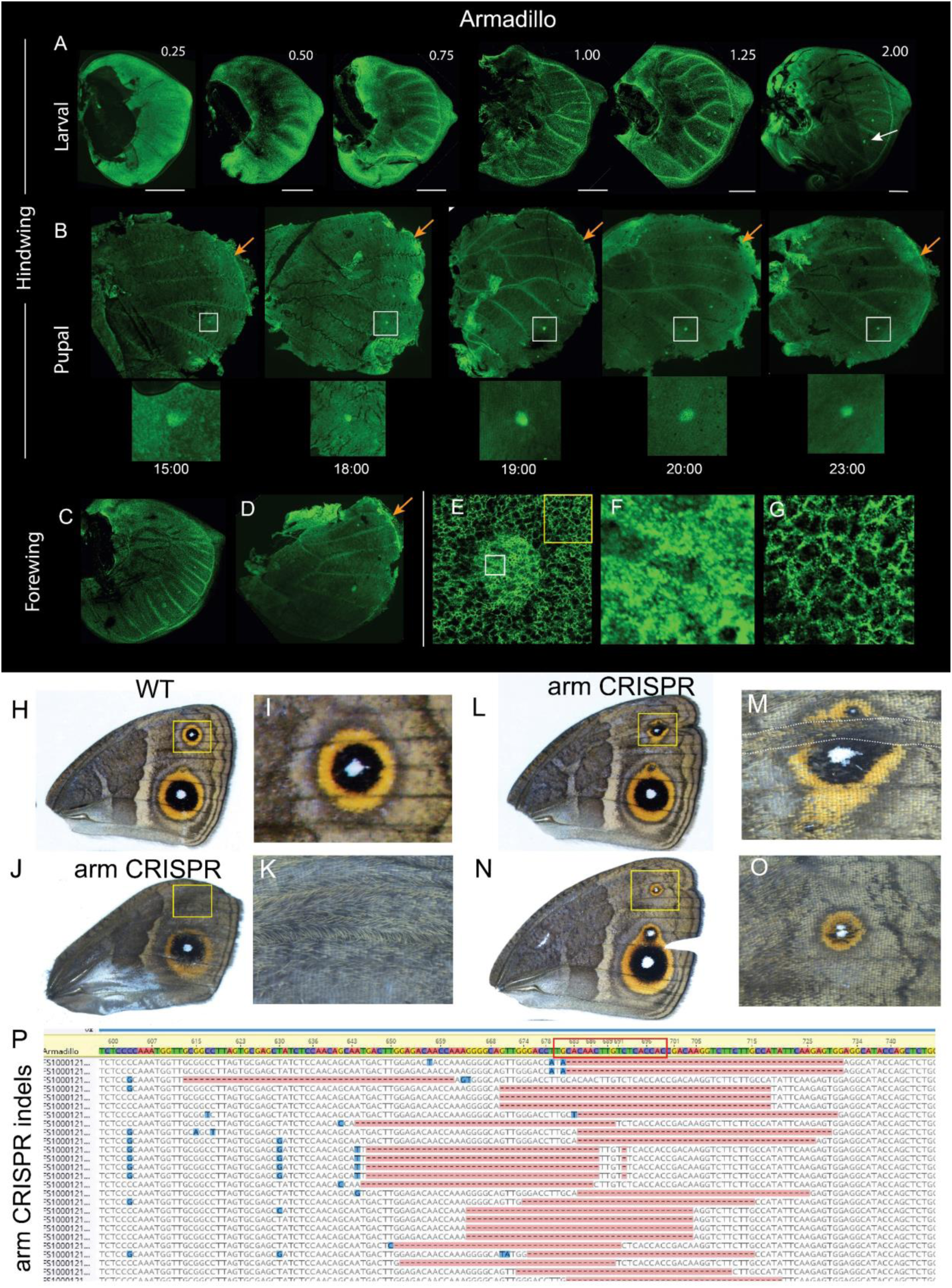
Localization and function of Armadillo (Arm) in *Bicyclus anynana*. (**A**) Arm is present homogeneously throughout the larval hindwing tissue till stage 0.75. At stage 0.75, eyespot focal expression is observed that continues throughout the larval stage together with expression along the wing margin and the wing veins (shown here till stage 2.00; for stage nomenclature please refer to Banerjee & Monteiro (2020)). Scale bar: 250 µm. (**B**) During pupal hindwing development, Arm is observed in the foci from 15-24 hrs after pupation. (**C**) Arm in larval forewings. (**D)** Arm in pupal forewings. (**E and F**) Arm is localized in the cytoplasm and in the nuclei of focal cells. (**G**) Outside the foci, Arm is present mostly at the cell membrane. (**H**) WT adult forewing. *arm* disruptions, using CRISPR-Cas9, resulted in the complete loss of an eyespot (**J, K**), or in two foci differentiating in a single wing sector (**L-O**). (**P**) Next-generation sequencing (NGS) of the affected wing region of the *arm* CRISPR individual in panels **J and K** showing deletions at the target site (red box).

### *arm* is essential for the differentiation of *B. anynana* eyespots

To identify the functional role of *arm* in eyespot development we injected embryos with Cas9 and a guide RNA targeting *arm*. Mosaic crispants showed either a complete loss of eyespots (one individual) (**Figure 1 J, K**), or split eyespots, where two foci differentiated side-by-side within a sector bordered by veins (**Figure 1 L-O**). The CRISPR phenotype was verified by Illumina paired-end sequencing where indels were observed at the CRISPR-Cas9 target site (**Figure 1P**). These results indicate that *arm* is essential for eyespot center differentiation and is likely involved in a previously proposed reaction-diffusion mechanism (see Discussion below).

### Canonical *wnts* (*wnt1, wnt6, and wnt10*) are expressed in larval wings and *wnt1* also in pupal wings

To localize all canonical *wnt* transcripts on the wing that could be responsible for the nuclear translocation of Arm in the eyespot focal cells (**Figure 1E, F**), we used hybridization chain reaction (HCR3.0) (Choi et al., 2018). *wnt1*, *wnt6*, and *wnt10* were all expressed along the wing margin during larval development (**Figure 2A; Figure S7A**), and there was no specific expression in the eyespot foci (**Figure 2A**). In 18-24 hrs pupal wings, however, we confirmed the expression of *wnt1* transcripts in the eyespot foci using *in-situ* hybridizations, as well as expression along the wing margin (**Figure 2B; Figure S7B**) (Özsu et al., 2017). During the pupal stage, the nuclear presence of Arm in the foci is likely driven by the locally transcribed *wnt1*. However, in the larval stage, nuclear Arm at the foci is probably driven by *wnt1*, *wnt6*, and *wnt10* produced along the wing margin, some distance away, and reaching the focal cells via some form of diffusion (or other form of transport), as previously modelled (Connahs et al. 2019).

**Figure 2.**
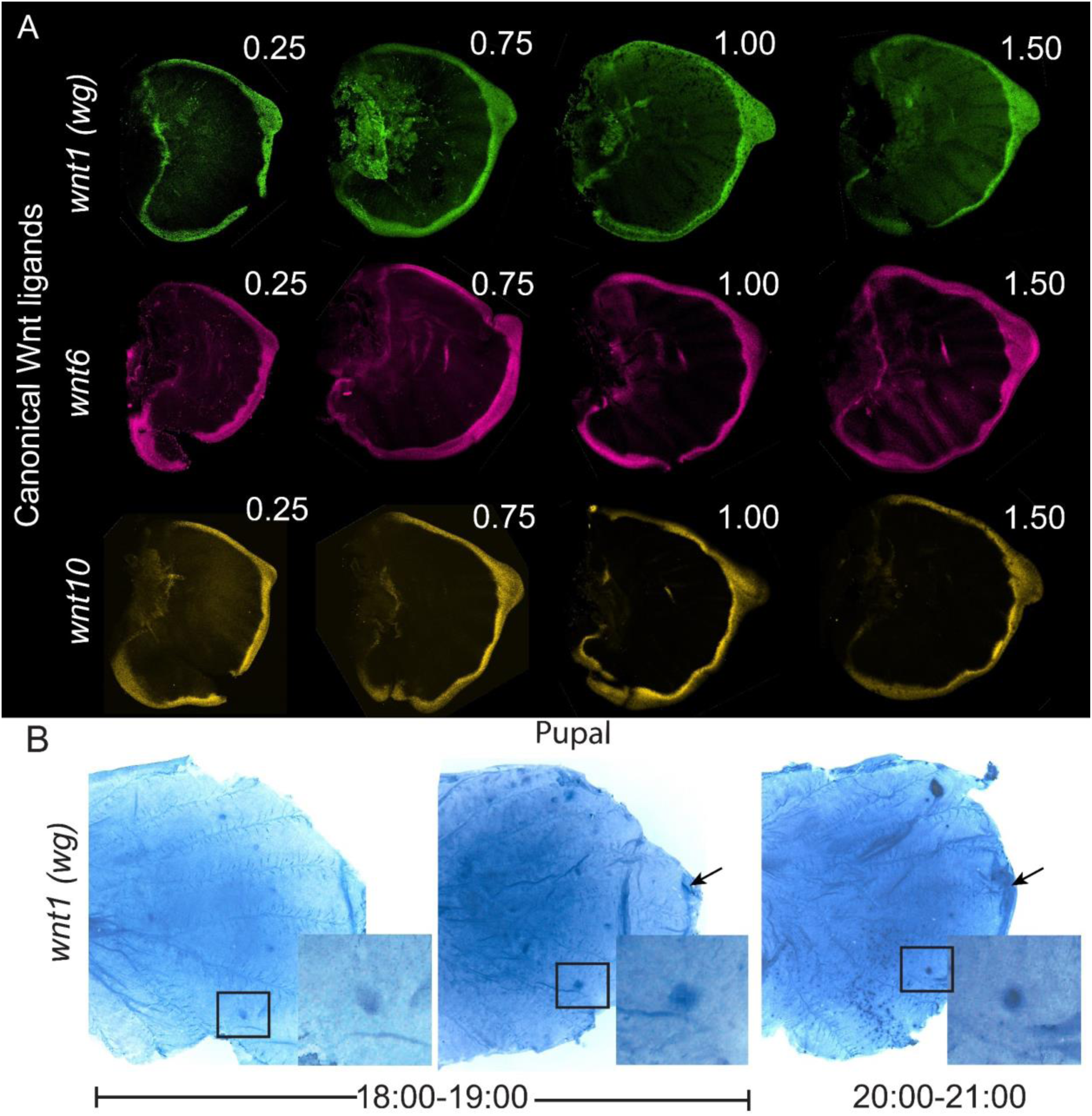
Expression of canonical *wnt* ligands using HCR and *in-situ* hybridization in larval and pupal wings. (**A**) The mRNA expression of three canonical *wnts*, *wnt1 (wg), wnt6,* and *wnt10,* is limited to the wing margin (larval stages 0.25-1.50) and there is no focal expression. (**B**) Expression of *wnt1* transcripts in the eyespot foci (black squares) and in the wing margin (black arrows) from 18 - 21 hrs of pupal wing development.

### Wnt1 target genes *Distal-less* (*Dll*) and *vestigial (vg*) are expressed in larval wings and *Dll* also in pupal wings

To investigate the domain over which Wnt1 glycoproteins might be activating target genes, we examined the co-expression of *wnt1* and two known targets of Wnt signaling, *Dll* and *vg*, in larval and pupal wings of *B. anynana*. In the *Drosophila* wing disc Wnt1 is secreted from the wing margin and travels into more interior wing regions where it activates *Dll* and *vg* at two different concentration thresholds (Neumann & Cohen, 1997). In the early larval wings of butterflies, we observed the expression of *Dll* in broad finger-like projections some cells away from the wing margin (**Figure 3B**), while *vg* is expressed in a broader domain (**Figure 3C**), consistent with *Drosophila* data (Neumann & Cohen, 1997). Interestingly, in a later larval stage (2.00) we observed *Dll* expression in the focal cells (**Figure 3E, F, I**) while *vg* was up-regulated in a slightly broader cluster of focal cells (**Figure 3G, H, J**). *wnt1*, however, was still not expressed in the foci at this stage (**Figure 2A, Figure S7A**). No expression of vg was observed in the pupal stage (**Figure S8**) likely because both Wnt and BMP signaling are involved in differentiating the eyespot rings (Banerjee et al., 2022), and their activity together activate *Dll,* but not *vg* (Diaz-Benjumea et al., 1994; Estella et al., 2008).

**Figure 3.**
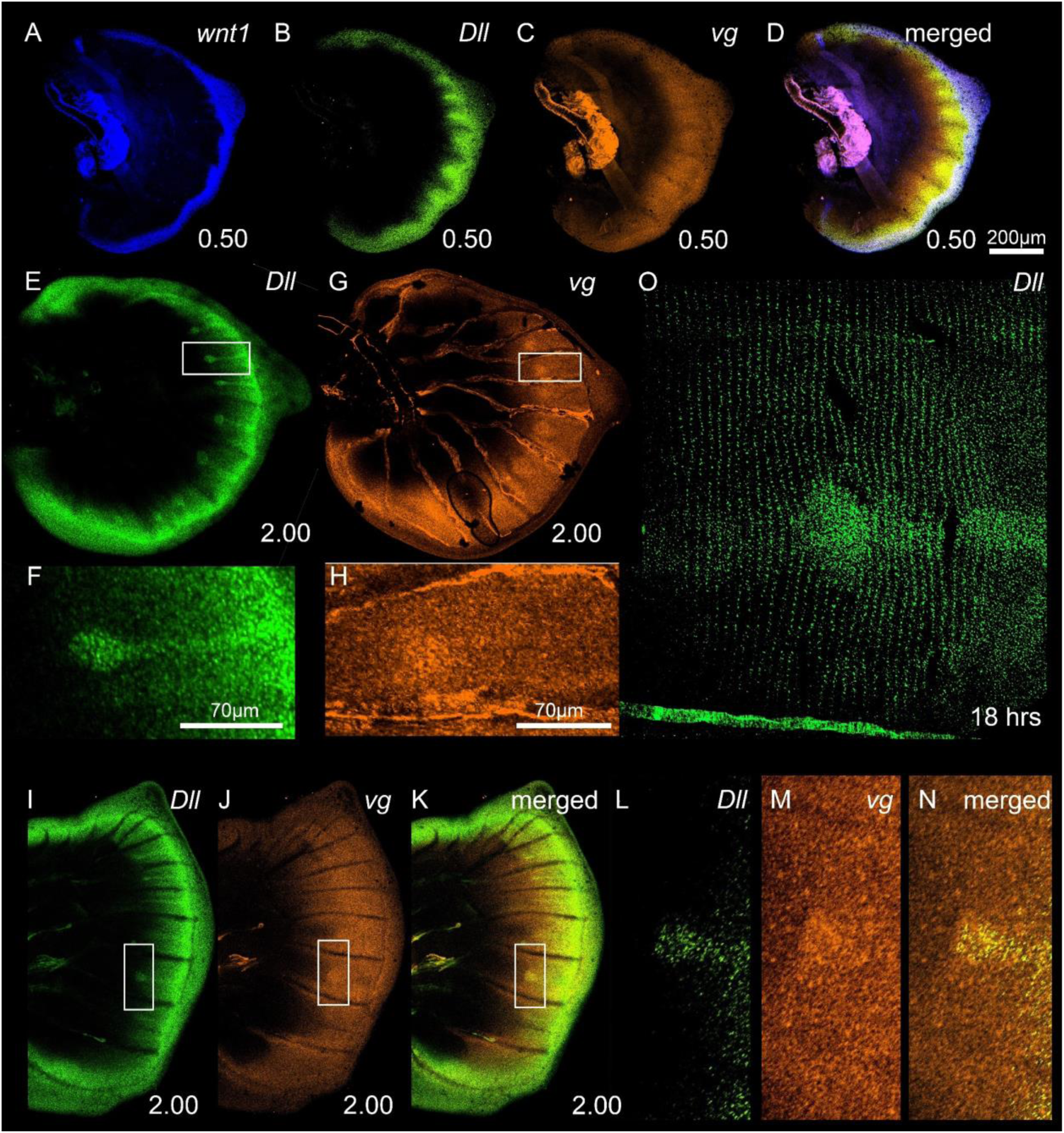
Expression of *wnt1, Dll,* and *vg* in larval wings. (**A**) *wnt1* is expressed along the wing margin. (**B**) *Dll* expression is observed some cells away from the wing margin, while (**C**) *vg* is expressed in a broader domain. (**D**) Merged expression of *wg, Dll,* and *vg* showing the range of a potential Wnt glycoprotein gradient activating its potential target genes. (**E, F**) Expression of *Dll* in an older larval wing (stage 2.00) in the foci and in the fingers from the wing margin. (**G, H**) Expression of *vg* showing a broader domain of expression both along the wing margin, and in the foci. (**I-N**) Co-expression of *Dll* and vg in the larval forewing showing expression in the eyespot center in a smaller domain for *Dll* and in a bigger domain for *vg*. (**O**) Expression of *Dll* in a 18 hr pupal wing showing expression in the future black scale cells.

### *wnt5, wnt7, wnt9*, and *wnt11* are not expressed in larval wings

To test whether *wnt5, wnt7, wnt9* and *wnt11,* typically associated with non-canonical Wnt signaling, could play a role in eyespot center differentiation we also examined their expression in larval wings. None of the genes showed any specific expression domain in the larval wings of *B. anynana* (**Figure 4A-H; Figure S9**). However, knockouts of *wnt7* using CRISPR-Cas9 resulted in ectopic veins and ectopic eyespots differentiating in the novel wing sectors (**Figure S12A-D**). Because ectopic veins often lead to the creation of additional wing sectors with eyespots, we cannot directly implicate *wnt7* in eyespot development. This gene appears to play a role, however, in vein development.

**Figure 4.**
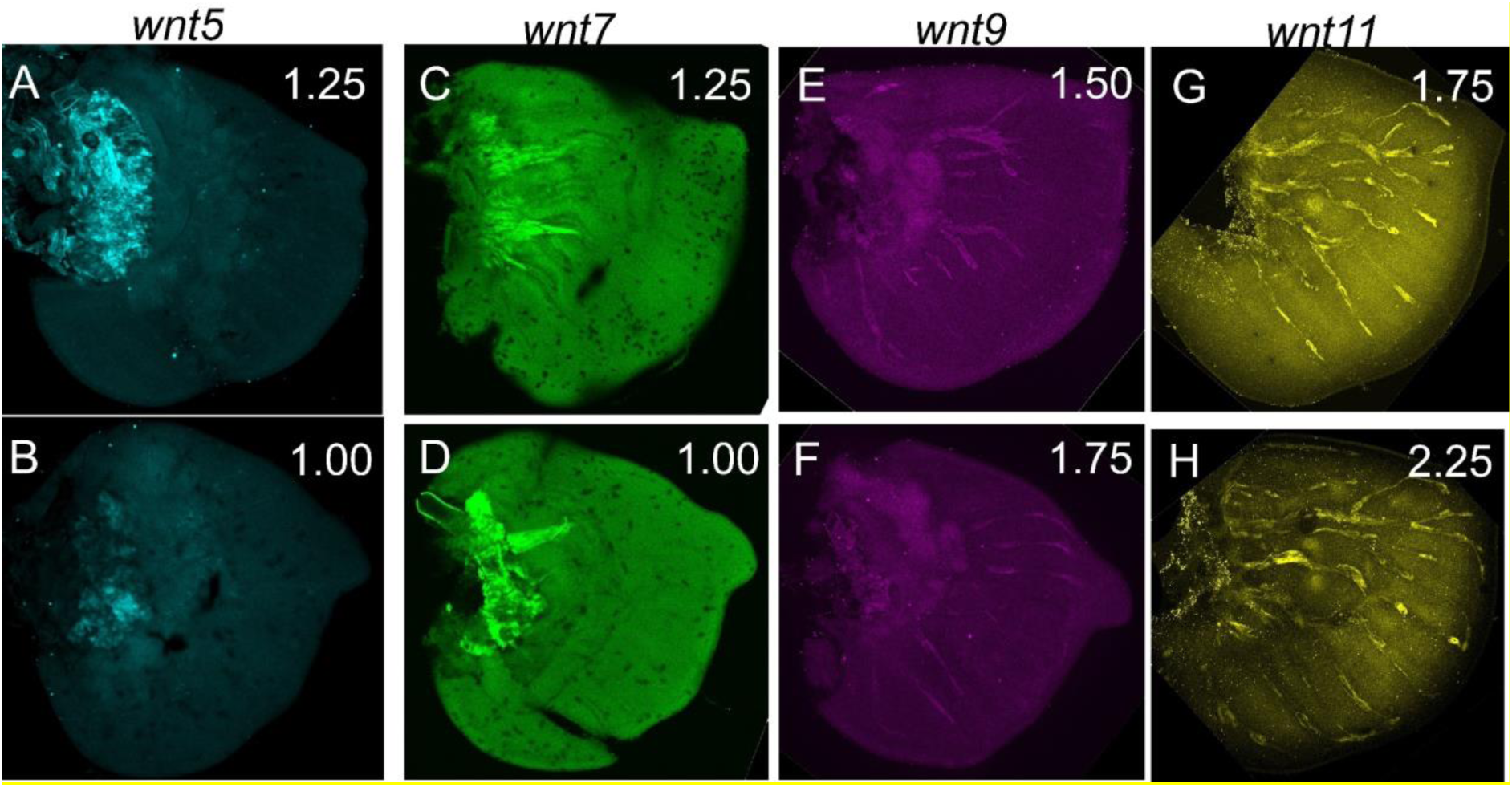
Expression of Wnt ligands *wnt5, wnt7, wnt9,* and *wnt11* in larval wings. No specific expression domain was observed for *wnt5* (**A, B**), *wnt7* (**C, D**), *wnt9* (**E, F**), and *wnt11* (**G, H**) during larval wing development in *B. anynana*.

### Expression of *frizzled4* and *frizzled9* in larval wings is anti co-localized

We next aimed to identify the Frizzled receptor(s) that might be used for Wnt signaling in larval wings. We first examined the expression of *frizzled4* and *frizzled9* transcripts in larval wings using HCR. In early stages (0.75-1.0) *frizzled4* was expressed homogeneously in the intervein cells (**Figure 5A, E; Figure S10A-B**), but at later stages (1.25-2.0) the expression was down-regulated in the eyespot foci and finger projections from the wing margin (**Figure 5H, K; Figure S10C-H**). The expression of *frizzled9* appeared complementary to that of *frizzled4* (**Figure 5C, G, J, and M**), with an early strong expression along the wing margin and in the finger-like projections at stage 0.75-1.25 (**Figure 5B, F, Figure S10Q, R**), that became restricted to the wing margin at later stages (Stage 2.00) (**Figure 5I, L; Figure S10M-T**). *frizzled4* also had lower expression in the lower posterior domain of late larval forewings where *frizzled9* had higher expression (**Figure 5H-J**).

**Figure 5.**
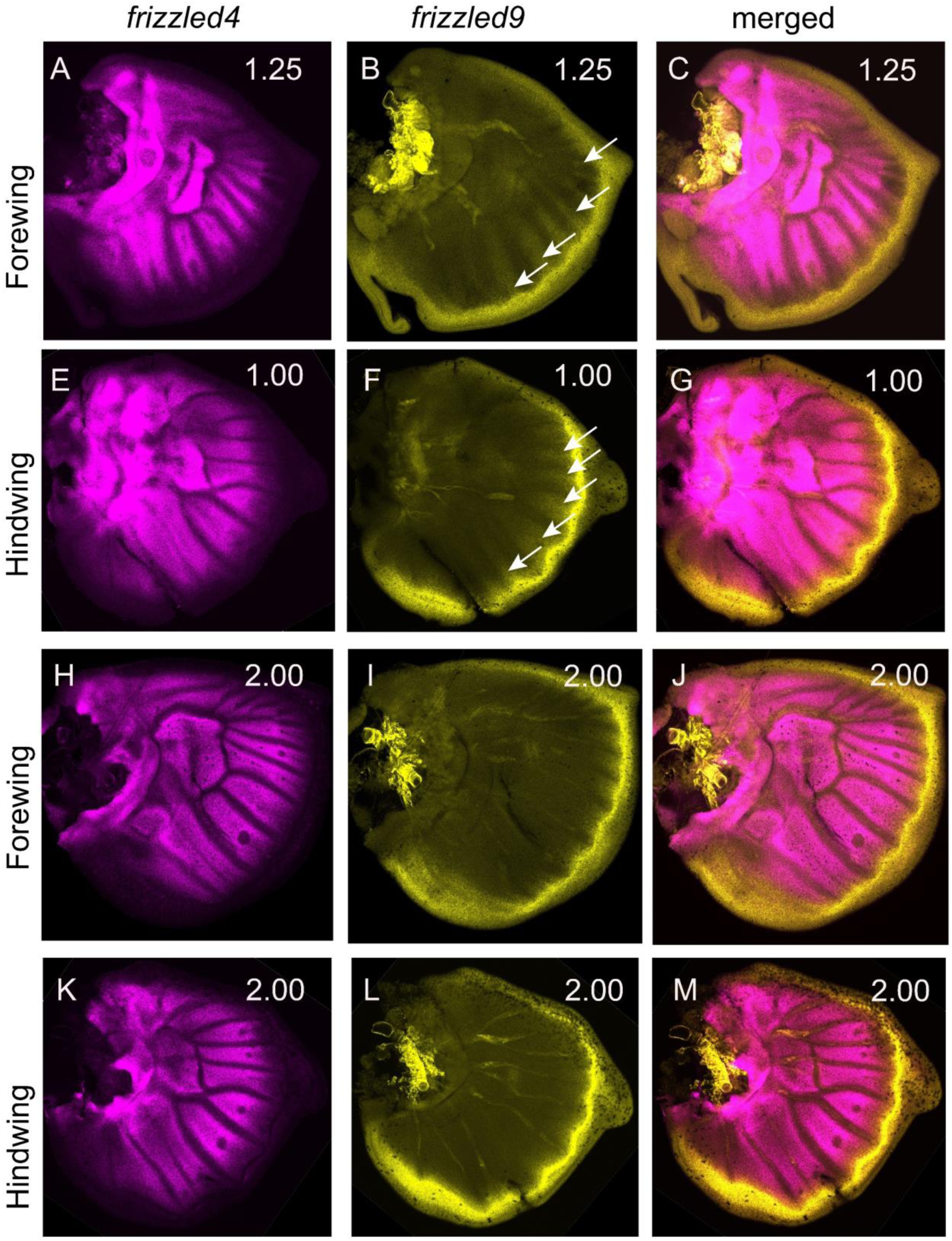
Expression of *frizzled4* and *frizzled9* in the larval wings of *B. anynana*. Expression of *frizzled4* and *frizzled9* in early larval forewings (**A, B**) and hindwings (**E, F**). **(C, G)** Merged channels of *frizzled4* and *frizzled9*. Expression of *frizzled4* and *frizzled9* in late larval forewings (**H, I**) and hindwings (**K, L**). (**J, M**) Merged channels of *frizzled4* and *frizzled9*.

### *frizzled2* is expressed in a proximal domain in larval wings and *frizzled* in eyespot foci in pupal wings

We performed HCR on two other potential receptors, *frizzled2* and *frizzled*, in both larval and pupal wings. In the larval stage, the expression of *frizzled2* was restricted to the proximal domain of both fore and hindwings (**Figure 6A-D**). *frizzled,* showed no specific expression in larval wings (**Figure 6E, F; Figure S11A-D**) but was strongly expressed in the eyespot foci in pupal wings at 18-24 hrs post pupation (**Figure 6G, H; Figure S11E-H**).

**Figure 6.**
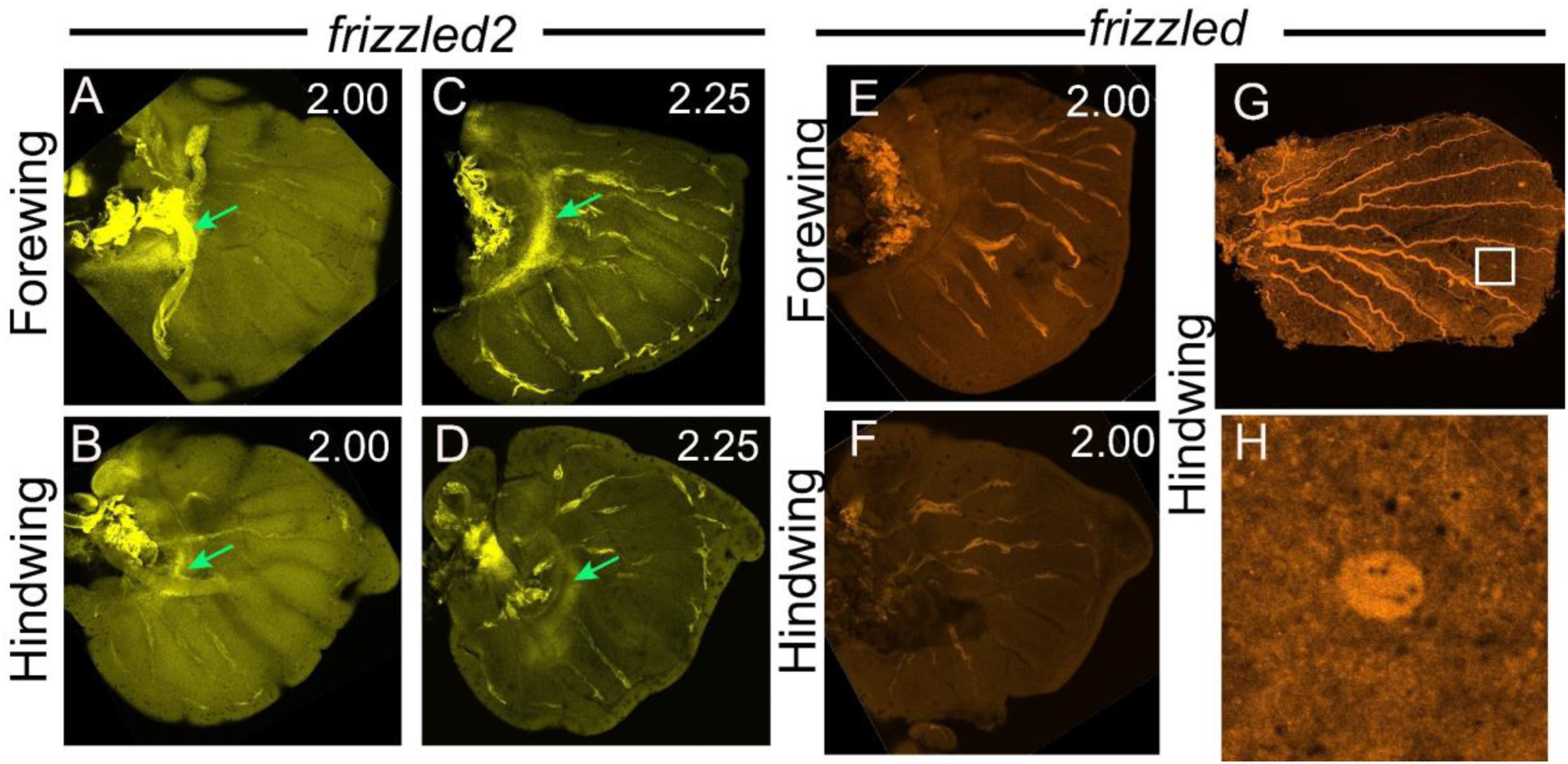
Expression of *frizzled2* and *frizzled* in larval and pupal wings. Expression of *frizzled2* in (**A-D**) larval wings along a proximal domain (green arrows). (**E**) There is no clear expression of *frizzled* in larval forewings and (**F**) hindwings. (**G, H**) Expression of *frizzled* in pupal eyespot foci in hindwings at 18-24 hrs.

### *frizzled2* is anti*-*colocalized with *frizzled4* and *frizzled9* in the eyespot field and pupal wing margin

Next, we investigated the 18-24hrs pupal wing expression of *frizzled2*, *frizzled4,* and *frizzled9*. During the pupal stage, *frizzled4* was newly expressed in the eyespot foci and in domains spanning the eyespot rings, bearing little resemblance to its early larval patterns (**Figure 7A, B; Figure S11K**). *frizzled2* was expressed in two large domains flanking each side of the central symmetry system, a proximal band domain, and a distal domain (**Figure 7C; Figure S11I-J**). In this distal domain, the expression of *frizzled2* was elevated in the eyespot foci but reduced in the areas of the future black and orange rings (**Figure 7C, D**). Co-expression of *frizzled2* and *frizzled4* showed the domain over which *frizzled4* was highly expressed had reduced levels of *frizzled2* (**Figure 7E-G; Figure S11L**). Lower levels of *frizzled 2* were also observed along the wing margin, where *frizzled9* was strongly expressed (**Figure 7H, I, and K**). The focal expression of *frizzled4* overlapped the expression of *wnt1* (**Figure 7L-N**).

**Figure 7.**
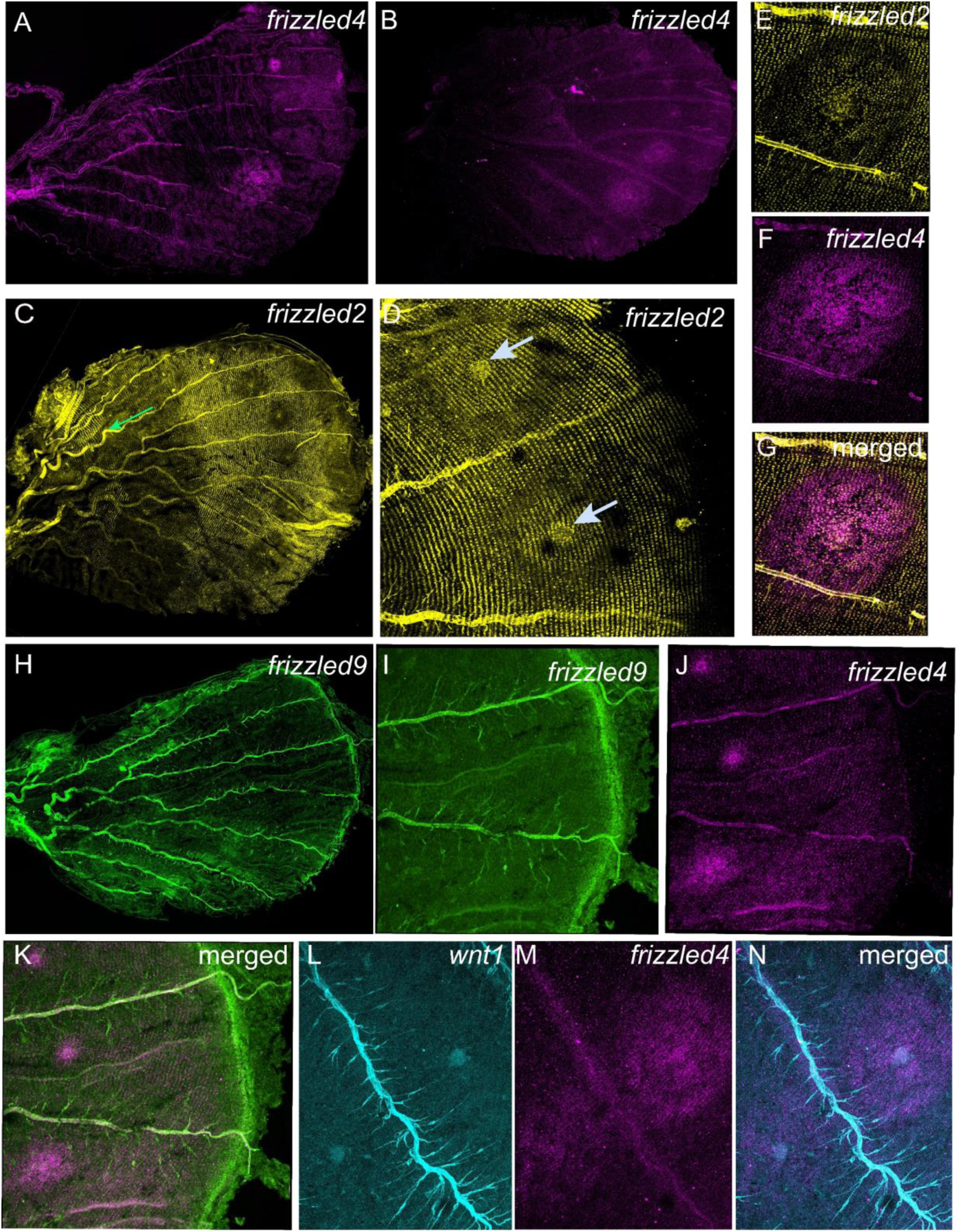
Expression of *frizzled2, frizzled4*, *frizzled9*, and *wnt1* in 18-24 hr pupal wings of *B. anynana*. Expression of *frizzled4* in the eyespot field of pupal forewings (**A**) and hindwings (**B**). (**C**) *frizzled2* is expressed in the proximal (green arrow) and distal domains straddling the central symmetry system of nymphalid butterflies (which consists of a central band running from the anterior to the posterior of the wing). In the distal domain, *frizzled2* has reduced expression in the eyespot field, and especially in the cells of the orange ring (**D**), but is present in the foci. Expression of (**E**) *frizzled2*, (**F**) *frizzled4*, and (**G**) merged channel in the forewing Cu1 eyespot. *frizzled2* is repressed in the domain where *frizzled4* is expressed. Pupal (**H**) hindwing showing the expression of *frizzled9* in the wing margin. Co-expression of (**I**) *frizzled9*, (**J**) *frizzled4*, and (**K**) merged. Expression of (**L**) *wnt1*, (**M**) *frizzled4*, and (**N**) merged expression in the hindwing.

### *frizzled4* functions in eyespot center differentiation and in planar cell polarity

To test the role of *frizzled4* we knocked it out using CRISPR-Cas9 at the embryonic stage. Some of the mosaic crispant wings differentiated two eyespot centers along the proximal distal axis (**Figure 8C-F; Figure S12E-G**). Other crispants also showed defects in the orientation of the black scale cells (**Figure 8G-I**). The knockout results were confirmed by sanger sequencing (**Figure 8J**).

**Figure 8.**
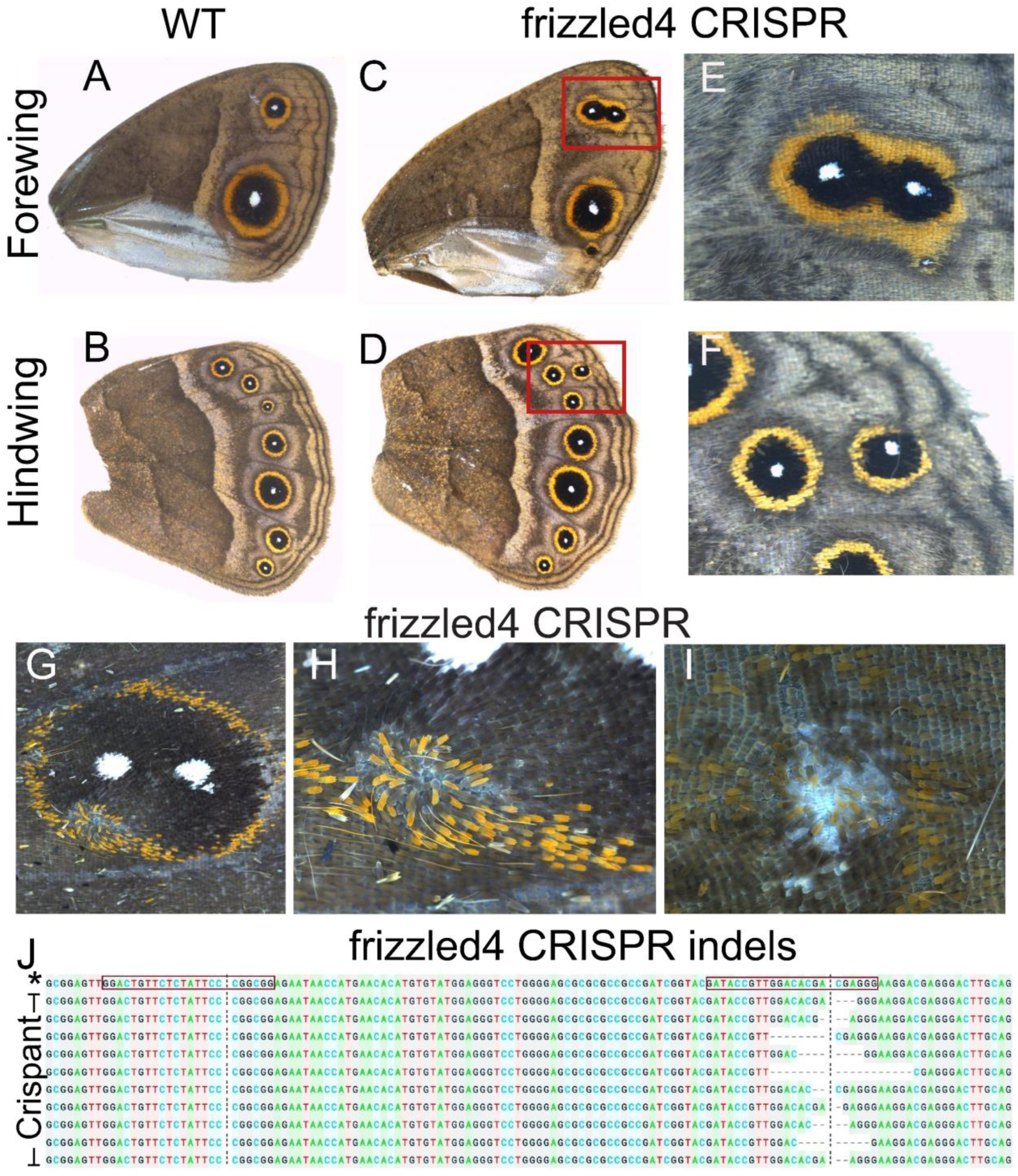
Function of *frizzled4* in eyespot center formation and planar cell polarity in *B. anynana.* (**A and B**) WT forewing and hindwing. (**C-F**) CRISPR-Cas9 on *frizzled4* resulted in the differentiation of two eyespots in the same wing sector. (**G-I**) CRISPR-Cas9 targeting *frizzled4* showed that *frizzled4* plays a role in planar cell polarity affecting scale orientation in the eyespot field. (**J**) Sequences from *frizzled4* CRISPR individuals showing indels at the second CRISPR target site. Target sites are highlighted in red boxes. Red boxes indicate the two target sequences. * Indicates the WT sequence.

## Discussion

### Canonical and non-canonical Wnt signaling are anti-localized on larval wings and necessary for the differentiation of eyespot centers

The differentiation of eyespot centers involves canonical Wnt signaling as shown by functional disruption of *arm*. In the larval stage, the canonical Wnt signaling transducer Arm gradually becomes present at high levels in the fingers, and in the nucleus of the eyespot centers, indicating the activation of Wnt signaling in those cells (**Figure 1E, F**). Disruptions of *arm* via CRISPR-cas9 resulted in either a split eyespot or no eyespots, confirming *arm*’s essential role in eyespot center differentiation (**Figure 1J-O**). However, none of the canonical *wnt* ligands tested, including *wnt1*, *wnt6* and *wnt10,* are present in the center of the eyespots at this stage (**Figure 2A**).

These results are consistent with the proposed mechanism of reaction-diffusion for eyespot center differentiation. This mechanism involves Wnt ligand production strictly at the wing margin, and Wnt glycoproteins diffusing from there into the more interior regions of the wing (Connahs et al., 2019). The reach of Wnt signal from the margin was visualized by the extent of the *vg* expression domain (**Figure 3C,G, and J**), a known response gene to marginal Wnt signaling in *Drosophila* wing discs (Neumann & Cohen, 1997). The proposed reaction-diffusion mechanism also involves the activation of a known second response gene, *Dll,* via Arm (Zecca et al., 1996). Previous work showed that when elongated clusters of *Dll* mutant cells (identifiable in those experiments due to scale color changes) were present in the middle of a wing sector, two eyespot centers differentiated side by side in that sector (Connahs et al., 2019). This is what we also observed in some Arm mutants (**Figure 1M and O**). These results, thus, mimic the *Dll* crispants and support the proposed reaction-diffusion model for eyespot center differentiation in the larval stage.

For Wnt ligands to reach the focal cells some distance away from the wing margin, they need to have a Wnt receptor expressed in those cells, and Frizzled9 is likely that receptor (**Figure 5B, F; Figure 9B**). This receptor is expressed not only along the wing margin but also in the intervein cells (**Figure 5B, F; Figure S10Q,R**) where Arm is also present (**Figure 1A, C**). In cell culture experiments Frizzled9 was shown to be involved in the activation of the canonical Wnt signaling and in the accumulation of Armadillo (β-catenin) in the cytoplasm (Karasawa et al., 2002). While functional experiments are still required, we propose that during larval wing development, canonical Wnt proteins are secreted from the wing margin and captured by Frizzled9 in the finger projections and eyespot foci. This results in accumulation of cytoplasmic Arm in those cells and eventually nuclear Arm (**Figure 1E, F**; **Figure 9B**). Frizzled9 is also most likely involved in the transduction of Wnt signaling to activate *Dll* and *vg* at different thresholds along the wing margin (**Figure 3; Figure 9B)** (Neumann & Cohen, 1997).

**Figure 9:**
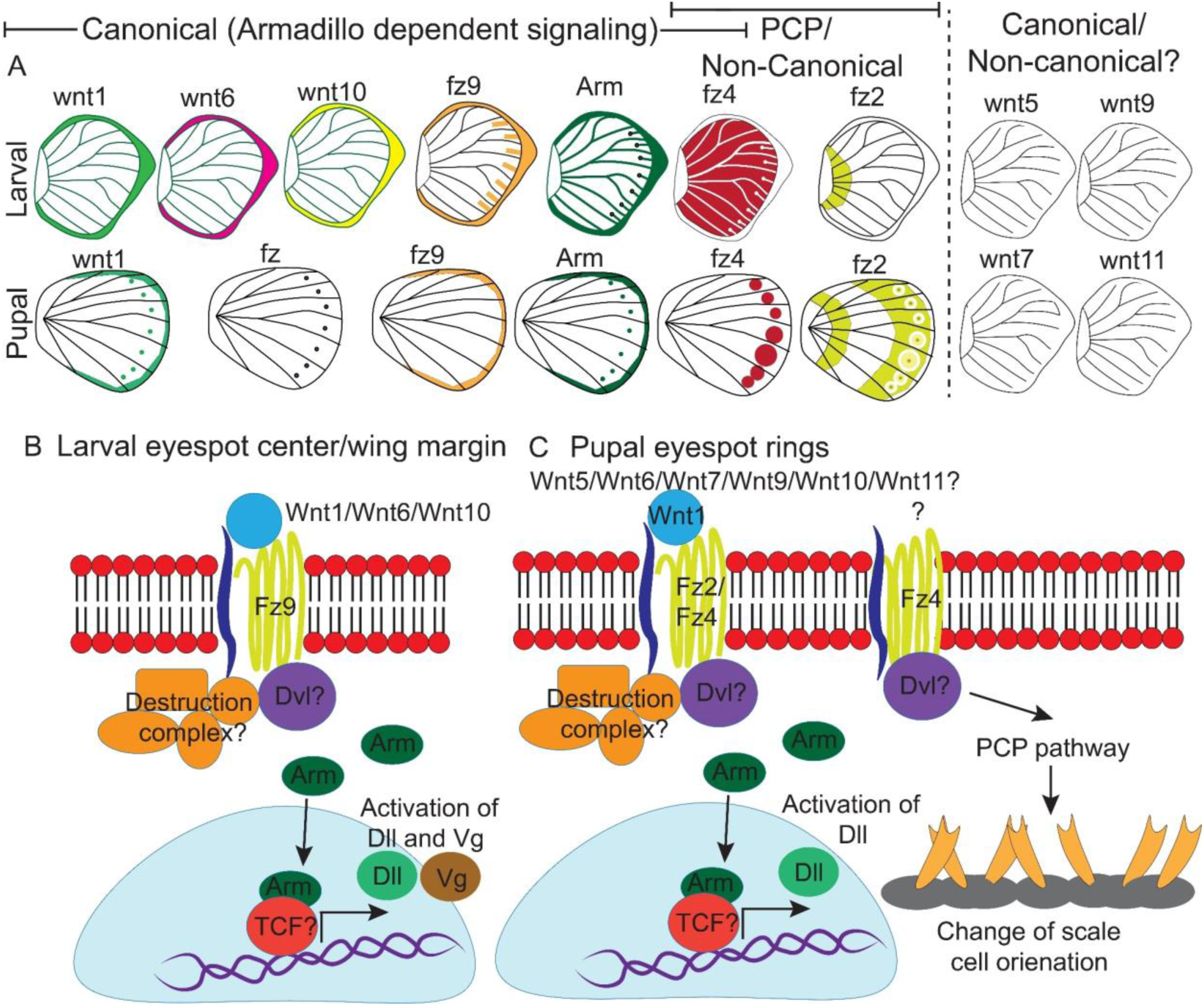
Expression of canonical and non-canonical Wnt ligands, receptors and signal transducer (Arm) in larval and pupal wings. (**A**) Expression patterns of *wnt1*, *wnt6*, and *wnt10*; receptors *frizzled2 (fz2)*, *frizzled9*(*fz9), frizzled (fz),* and *frizzled4* (*fz4)*; and the signal transducer Arm in larval and pupal wings. No expression pattern is observed for the Wnt ligands *wnt5*, *wnt7, wnt9*, and *wnt11*. (**B**) During larval wing development, ligands of canonical Wnt signaling bind to the receptor Fz9 preventing the proteasomal degradation of Arm and promoting its accumulation in the nucleus of cells in the foci and wing margin. Wnt signaling activates target genes *Dll* and *vg*. (**C**) In the pupal wing, Wnt1 is a likely morphogen binding to receptors Fz2 and Fz4 and leading to canonical Wnt signaling. Here the transcription factor *Dll* is likely responding to this signal. *Dll* is responsible for the differentiation of black scales in the eyespot (Connahs et al., 2019). The expression and function of *fz4* in the eyespot field indicates active planar cell polarity signaling in those cells that keeps scales oriented in the correct way on the wing.

The other *frizzled* genes had either no wing expression (*frizzled*), proximal wing expression (*frizzled2*), or anti-colocalized expression with *frizzled9*/Arm (*frizzled4*). The presence of *frizzled2* in the proximal domain of larval wings is likely due to the repression of *frizzled2* by Wnt signaling from the wing margin as previously shown in *Drosophila* larval wings (Cadigan et al., 1998).

The differentiation of eyespot centers also involves non-canonical Wnt signaling, which may cross-regulate canonical Wnt signaling. *frizzled4,* a receptor typically associated with non-canonical Wnt signaling (Descamps et al., 2012) is anti-colocalized with Arm in the wing margin, veins, eyespot foci and finger projections (**Figure 9A; Figure 5A, E, H, and K; Figure S10A-H**), all the areas where Arm is present (**Figure 1A, C**). None of the proposed non-canonical Wnt ligands (*wnt5, wnt7*, *wnt9,* and *wnt11*), however, are co-expressed with *frizzled4* (**Figure 4; Figure 9A; Figure S9**), indicating that non-canonical Wnt signaling is likely transduced through Frizzled4 independently of any Wnt ligand, similar to what has been proposed for *Drosophila* wings (Ewen-Campen et al., 2020; Yu et al., 2020). Disruptions of *frizzled4* in *B. anynana* resulted in the appearance of two eyespots along the proximo-distal axis of a wing sector (**Figure 8C-F; Figure S12E-G**), which phenocopies a *Dll* over-expression phenotype previously studied and modeled via a reaction-diffusion mechanism using canonical Wnt signaling (Connahs et al. 2019). The loss of *frizzled4* likely led to the over-expression of canonical Wnt signaling and over-expression of target genes such as *Dll* (and *vg*). *frizzled4*-mediated signaling has been proposed to repress canonical Wnt signaling in arterial network organization in mice (Descamps et al., 2012). We propose, thus, that both canonical and non-canonical Wnt signaling pathways are interacting with each other, and are both essential for eyespot center differentiation.

### *frizzled4* is involved in regulating scale polarity in the eyespot field

*frizzled4* appears to also be involved in the orientation of the scale cells in the eyespot rings of *B. anynana*. The gene is expressed in the eyespot field during the pupal stage (**Figure 7A,B; Figure 9A**), and disruptions of *frizzled4* resulted in scale orientation defects (**Figure 8G-I; Figure 9C**). The planar cell polarity (PCP) pathway, however, is likely independent of any Wnt ligand as previously shown in *Drosophila* wings (Ewen-Campen et al., 2020; Yu et al., 2020). This pathway, when disrupted, results in changes in the orientation of cellular protrusions such as trichomes in *Drosophila* wings (Ewen-Campen et al., 2020) and mouse skin (Guo et al., 2004). It appears that scales are behaving in the same way as trichomes in *Drosophila* wing, despite not being homologous traits.

### Wnt1 is likely working with Frizzled2 and Frizzled4 receptors and the transcription factor Arm to determine eyespot size during the pupal stage

Wnt1 was previously proposed as a morphogen produced in the eyespot centers (Monteiro et al., 2006) to regulate eyespot size in *B. anynana* (Özsu et al., 2017). The mechanism by which Wnt1 operates in the determination of the eyespot rings, however, had not yet been explored. In the present work we confirmed the presence of *wnt1* mRNA in the eyespot central cells during early pupal wing development (Özsu et al., 2017)(**Figure 2B, Figure S7B**). The Wnt1 ligand is likely being secreted by these central cells and captured by the receptors Frizzled4 and Frizzled2, whose transcripts are being expressed across the whole eyespot field (**Figure 7A-D; Figure S11I-L; Figure 9C**). Active canonical Wnt signaling via Frizzled2 should result in higher levels of Arm in the eyespot rings. We, however, only observed strong Arm expression in the center of the eyespot during pupal wing development (**Figure 1B**). The expression of Arm in the central cells is likely due to the co-expression of *frizzled2* and *frizzled* (**Figure 7E-G; Figure S11L; Figure 9A**,) during pupal wing development. In the other ring cells Wnt signaling is probably being transduced in a non-canonical fashion.

### *frizzled4* likely functions in canonical and non-canonical signaling based on the developmental stage

In *B. anynana,* Frizzled4 likely functions in activating both canonical and non-canonical Wnt signaling. During the larval stage, Frizzled4 is likely transducing non-canonical Wnt signaling across most of the wing, while during the pupal stage it is transducing both canonical and non-canonical Wnt signaling. During the larval stage, *frizzled4* expression anti-colocalizes with Arm expression (**Figure 1A, C; Figure 5A,E,H,K; Figure 9A**) and is likely working in non-canonical Wnt signaling. During the pupal stage, *frizzled4* expression overlaps that of *wnt1* (**Figure 7L-N**) and Arm in the eyespot centers (**Figure 1B**) and might be working with these other genes to regulate eyespot size (**Figure S12E, F**) via canonical Wnt signaling (**Figure 9C**). In addition, *frizzled4* functions in the pupal stage to maintain polarity of the scale cells in eyespots (**Figure 8G-I; Figure 9C**). Frizzled4 has been shown to function in both the canonical (Tickenbrock et al., 2008) and non-canonical PCP pathway (Descamps et al., 2012), depending on the type of Wnt co-receptor (arrow/LRP) and ligand present in each context.

### Spatial regulation of *frizzled2 and frizzled 4* in the eyespot field

During the first 18-24 hrs of pupal wing development we observe a very precise control of the domains over which *frizzled2* and *frizzled4* are expressed (**Figure 7**), suggesting they are regulated (and cross-regulated) by the signaling pathways where they function. Both *frizzled2* and *frizzled4* are expressed in the eyespot focal cells, but *frizzled2* is anti-colocalized with *frizzled4* in the rest of the eyespot field (**Figure 7E-G; Figure S11L**).

Similarly, cross-regulation might be happening between a Wnt signaling pathway using *wnt1*, *frizzled9*, and Arm, expressed along the wing margin (**Figure 1B, Figure 2B, Figure 7H**) and *frizzled 2*, expressed at lower levels in the margin and chevron area (**Figure 7C**).

We propose a model (**Figure 10**) for the spatial regulation of *frizzled2* which involves down-regulation of this gene by Wnt signaling from the margin and from the eyespot centers. *frizzled2* is initially homogeneously expressed in the pupal wing cells, but both Wnt signaling from the wing margin, involving Wnt1, Frizzled9, and Arm, and Wnt signaling from the eyespot center, involving Wnt1, Frizzled4, and Arm, repress the activity of *frizzled2* in the respective domains resulting in two disjointed, but precise domains of *frizzled2* expression (**Figure 10**). This model is consistent with Wnt1-dependent signaling in *Drosophila* repressing *frizzled2* in the dorsal-ventral axis of the wing disc and in the segments of the embryo (Cadigan et al., 1998; Lecourtois et al., 2001).

**Figure 10.**
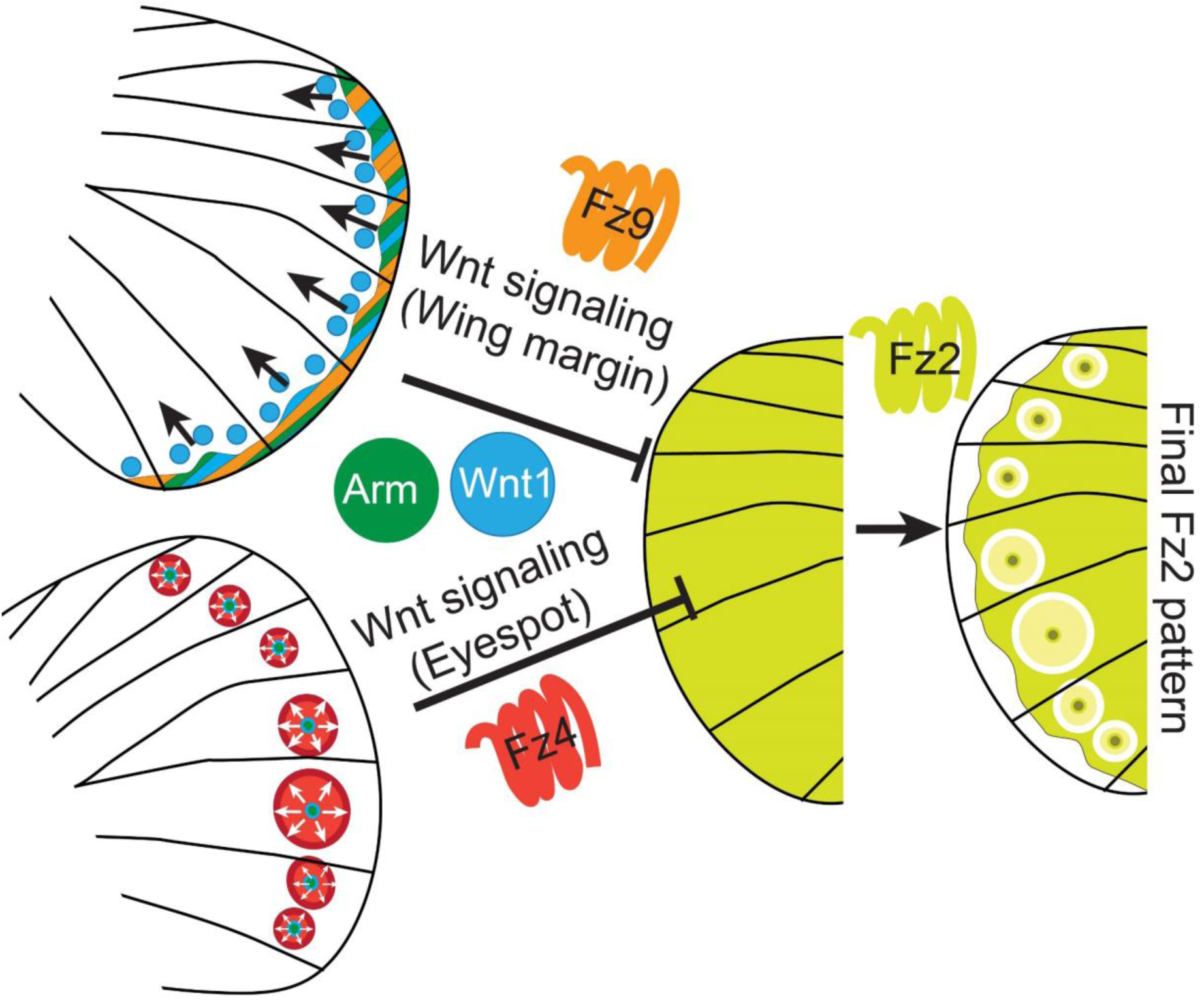
A model for the spatial control of *frizzled2* expression. In this model *frizzled2,* initially homogeneously expressed in the pupal wing tissue, becomes repressed in two distinct domains via Wnt signaling from the wing margin and from the eyespot central domain cells.

In summary, we illustrated the expression and function of different Wnt signaling pathway members in larval and pupal wings of *B. anynana* wings that contribute to the precise differentiation of the eyespot color pattern. Further studies are necessary to examine the function of several of these genes, especially that of *frizzled9*, and the interaction between *frizzled4* and *frizzled2*. The interaction of Wnt signaling with BMP also needs further study, as this has been proposed to function in the reaction diffusion mechanism proposed for eyespot center formation during larval stage (Connahs et al., 2019), and ring differentiation during the pupal stage (Banerjee et al., 2022). The Wnt signaling mechanism, however, is likely conserved in other nymphalid butterfly species with eyespots, as observed with the expression pattern of Arm (**Figure S6**) and canonical Wnt ligands (Martin & Reed, 2014).

Studying complex signaling pathways, such as Wnt signaling, in simple 2D traits such as butterfly eyespots may help unravel the function of this pathway in more complex 3D traits such as legs (Diaz-Benjumea et al., 1994; Estella et al., 2008), antennae (Murugeshan et al., 2022), and horns (Hu et al., 2019; Wasik & Moczek, 2012), where Wnt signaling is also fundamental to trait development.

## Methods

### Sequence alignment and phylogenetic tree construction

Nucleotide sequences were obtained from ncbi and flybase. Alignments were carried out using ClustalW (Thompson et al., 1994) with the default parameters in ‘SLOW/ACCURATE’ alignment option in GenomeNet.

Using RAxML(Randomly Accelerated Maximum Likelihood) v8.1.20 a maximum likelihood tree was created with model PROTGAMMAJTT and default parameters with 100 bootstraps (Stamatakis, 2014) using ETE v3.1.1 (Huerta-Cepas et al., 2016) implemented on GenomeNet. Similar trees were obtained with Fasttree with slow NNI and MLACC = 3 (to make the maximum-likelihood NNIs more exhaustive) (Price et al., 2009) ETE v3.1.1 (Huerta-Cepas et al., 2016).

### Rearing Bicyclus anynana

*Bicyclus anynana* larvae and adults were fed corn leaves and mashed bananas, respectively. The individuals were reared at 27°C with a 12-12 hrs day-night cycle under 60% humidity.

### CRISPR-Cas9

*arm, wnt7,* and *fz4* embryonic CRISPRs were carried out based on the protocol described in Banerjee & Monteiro (2018) and Murugesan et al (2022). One to two RNA guides were designed using CRISPRdirect or CHOPCHOP to target the coding sequence of these genes (see supplementary file for sequences). Cas9 protein (IDT, Cat No. 1081058) (300ng/μl) along with the guide RNA at a concentration of 300 ng/μl were mixed in molecular grade water with Cas9 buffer (NEB, Cat No. M0386S). Embryos were injected 6 hrs after egg laying for *arm* and 3hrs for *frizzled4* and *wnt7*. Embryos were kept in Petri dishes inside a temperature-controlled incubator with moistened cotton to maintain humidity. Larvae were reared in mesh cages and fed young corn leaves. After pupation, the larvae were transferred to plastic containers, and after eclosion the adults were frozen at -20°C and imaged under a Leica DMS1000 microscope.

For NGS sequencing, DNA was extracted from affected wings using an Omega tissue DNA extraction kit (Cat No. D3396-01). For the *arm* crispants, primers flanking 276 bps around the CRISPR target site were used to amplify the region of interest (primer sequences in **Table S1**). Adapters and indices were added to the PCR product via a two-step PCR reaction (**Table S1**). Followed by purification of the PCR products. The samples were sequenced using an Illumina iSeq 100 sequencer. Reads were aligned to the reference WT arm sequence using Geneious R10 (Kearse et al., 2012).

For Sanger sequencing of the crispants and wild-type, DNA was extracted as described above using Omega tissue DNA extraction kit. PCR amplified using the gene-specific primers (primer sequences in Table S1) and purified. Sequencing was carried out at 1^st^ Base, Singapore. CRISPR indel was analysed using Synthego Performance Analysis, ICE Analysis. 2019. v3.0. (last accessed Oct 2022) (https://www.synthego.com/products/bioinformatics/crispr-analysis).

### Immunostainings

Moment of pupation was timed using an Olympus tough TG-6 camera and pupal wings were dissected and stained based on a protocol described in Banerjee & Monteiro (2020b). Briefly, wings were dissected in 1x PBS under ice for larval wings and in 1x PBS at room temperature for pupal wings. Afterward, wings were transferred to fix buffer with 4% formaldehyde in ice for larval wings and in room temperature for pupal wings. After fixation, the wings were transferred to ice, washed with 1x PBS, and kept in block buffer overnight. Primary antibodies were diluted in wash buffer at the concentration of 1:1000 anti-Arm (Rat; Banerjee & Monteiro 2020b) and 1:20000 anti-Spalt (Guinea Pig GP66.1; Oliver et al., 2012) and stained overnight at 4°C. Afterwards, the wings were washed four times in wash buffer, for 15 mins each time. Wings were then incubated in 1:500 secondary antibody at the concentration 1:500 with anti-rat AF488 (Invitrogen, #A-11006) and anti-Guinea pig AF555 (Invitrogen, # A-21435), followed by four washes in wash buffer. Finally, the wings were mounted in an inhouse mounting media (see Supp table. Sx) and imaged under an Olympus fv3000 microscope.

### Enzyme based in-situ hybridization

In-situ hybridization experiments were carried out based on the protocol described in Banerjee et al. (2020). Pupal wings were dissected in 1x PBS and fixed in 4% formaldehyde in 1x PBST at room temperature. After fixation wings were treated with proteinase k for five mins and afterwards with 100 mg/ml glycine in 1x PBST. Wings were then washed with 1x PBST and transferred to pre-hyb buffer at 65°C followed by incubation in hybridization buffer with probes against *wnt1* (see suppl file for sequence) at 65°C for 16-24 hrs. After hybridization, wings were washed with pre-hybridization buffer at 65°C, five times, for 30 mins each time. Afterwards, wings were brought down to room temperature and washed four times with 1x PBST. Wings were then incubated in block buffer for 60 mins, followed by incubation in anti-digoxygenin (Sigma-Aldrich, Cat No. 11093274910) at the concentration of 1:2000 in block buffer. Wings were then washed five times with block buffer, 10 mins each time. Wings were then washed two times in alkaline-phosphatase buffer, for five mins each time. After washing, wings were transferred to alkaline-phosphatase buffer supplemented with NBT-BCIP (Promega, Cat No. S3771) and left for 4-6 hrs in the dark for color to develop. Once the color developed, wings were washed two times with 1x PBST and mounted in 60% glycerol and imaged under a Leica DMS1000 microscope.

### Fluorescent based in-situ hybridization (HCR3.0)

Fluorescent in-situ hybridization was performed based on the protocol developed by Choi et al., 2018 with few modifications optimized for butterfly wing tissue. Briefly wings were dissected in 1x PBS and fixed in 4% formaldehyde in 1x PBST. After fixation wings were washed with 1x PBST and 5x SSCT and permeabilized using a detergent solution (Bruce et al., 2021). Wings were again washed with 5x SSCT followed by addition of 30% probe hybridization buffer. Afterwards wings were incubated overnight at 37°C in a chamber with 30% probe hybridization buffer and HCR probes. The next day wings were washed with five times with 30% probe wash buffer at 37°C followed by two washes with 5x SSCT at room temperature. Wings were then incubated in amplification buffer with secondary fluorescent hairpin probes in the dark for 16-20 hrs. The next day wings were washed twice in 5x SSCT, mounted in an inhouse mounting media, and imaged under an Olympus fv3000 microscope.

## Author’s contribution

Conceptualization: T.D.B., A.M.; Methodology and investigation: T.D.B., S.N.M; Validation: T.D.B.; Formal analysis: T.D.B., A.M.; Resources: A.M.; Data curation: T.D.B.; Writing - original draft: T.D.B.; Writing - review & editing: T.D.B, A.M.; Visualization: T.D.B.; Supervision: A.M.; Project administration: A.M.; Funding acquisition: A.M.

## Acknowledgement

We would like to thank Dr. Eunyoung Chae’ lab and Yi Yun for help with the sequencing of the armadillo CRISPR individuals. We would like to thank the imaging facility at CBIS, NUS for access to the Olympus fv3000 microscope, and Anupama Prakash for helping with *frizzled4* crispants images.

## Supplementary information

**Figure S1:**
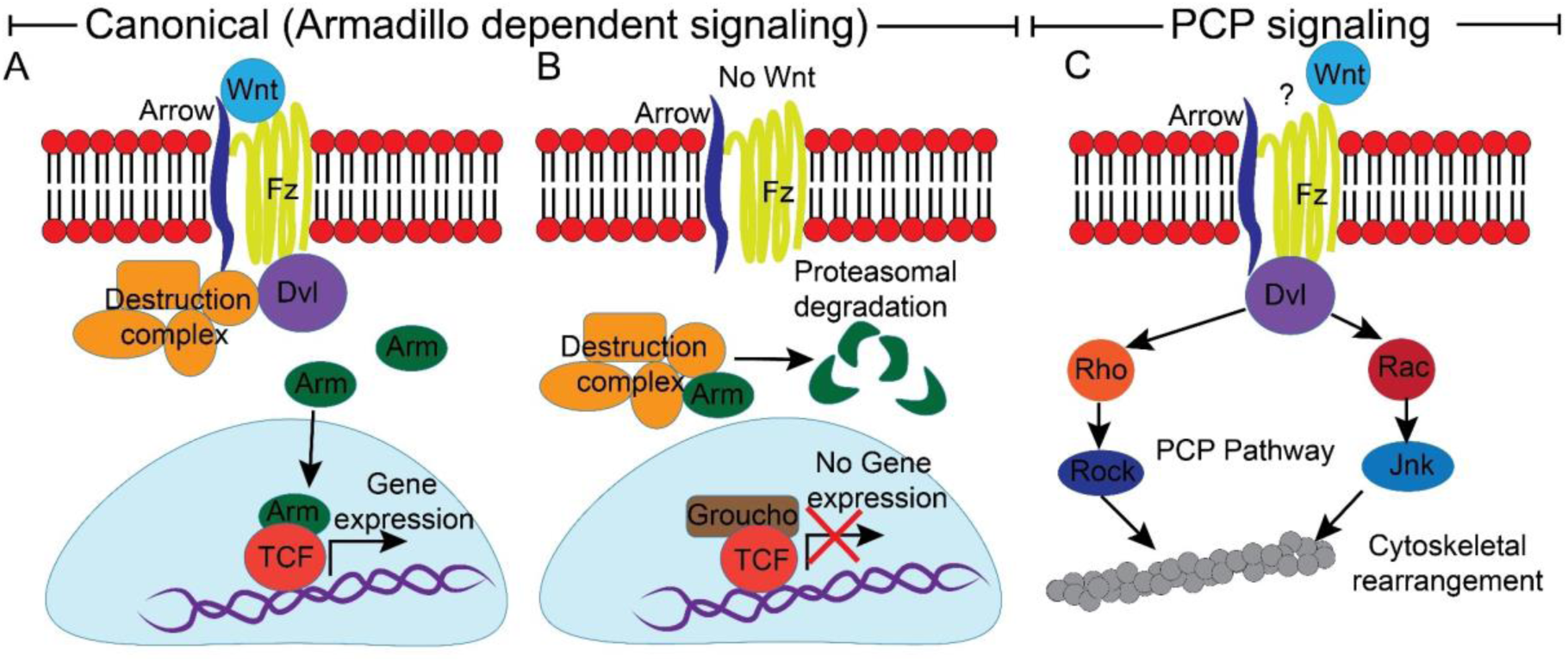
Canonical (Armadillo dependent) and PCP (non-canonical) Wnt signaling. (**A and B**) In the canonical Wnt signaling in the presence of Wnt the receptor Frizzled (Fz) and co-receptor Arrow with the help of Dishevelled (Dvl) prevents the destruction complex from destroying cytoplasmic Arm. Arm moves into the nucleus and with the help of co-transcriptional activator TCF activates gene expression. In the absence of Wnt, the destruction complex helps in the proteasomal degradation of Arm, and due to the absence of Arm in the nucleus the Wnt repressor Groucho binds TCF and prevents gene expression. (**C**) In the non-canonical planar cell polarity pathway Dvl activates the G-protein complex Rho and Rac which activates the Rock and Jnk signaling respectively. This signaling pathway is responsible for the cytoskeletal rearrangement of the cell.

**Figure S2.**
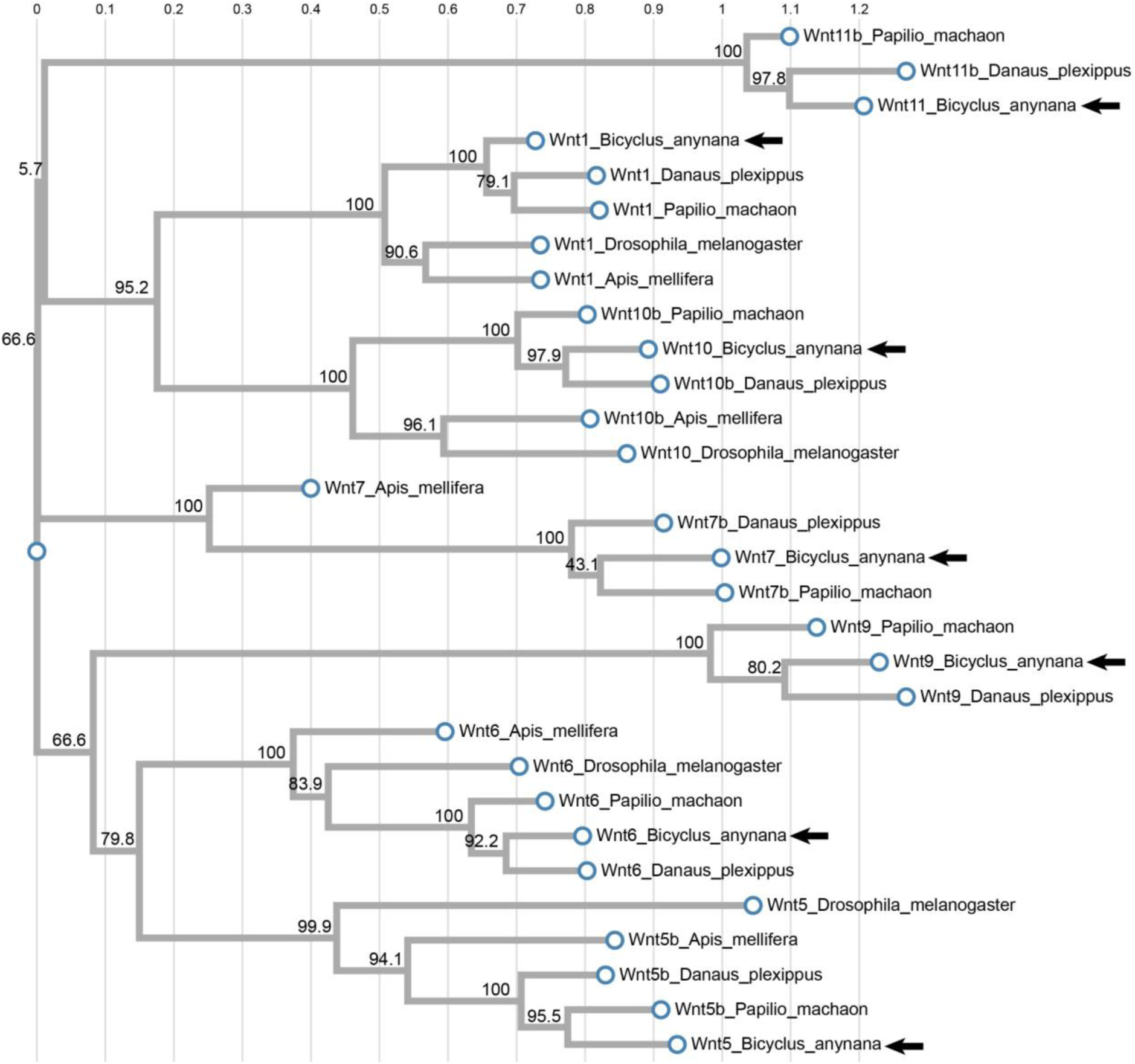
Phylogenetic tree of *wnt* genes created using fasttree (Maximum likelihood). The tree clusters the Wnt family genes together. Vertical lines indicate mean number of nucleotide substitutions per site. Numbers above branches represents bootstrap branch support. Arrows point to the seven *wnt* genes in the genome of *B. anynana*.

**Figure S3.**
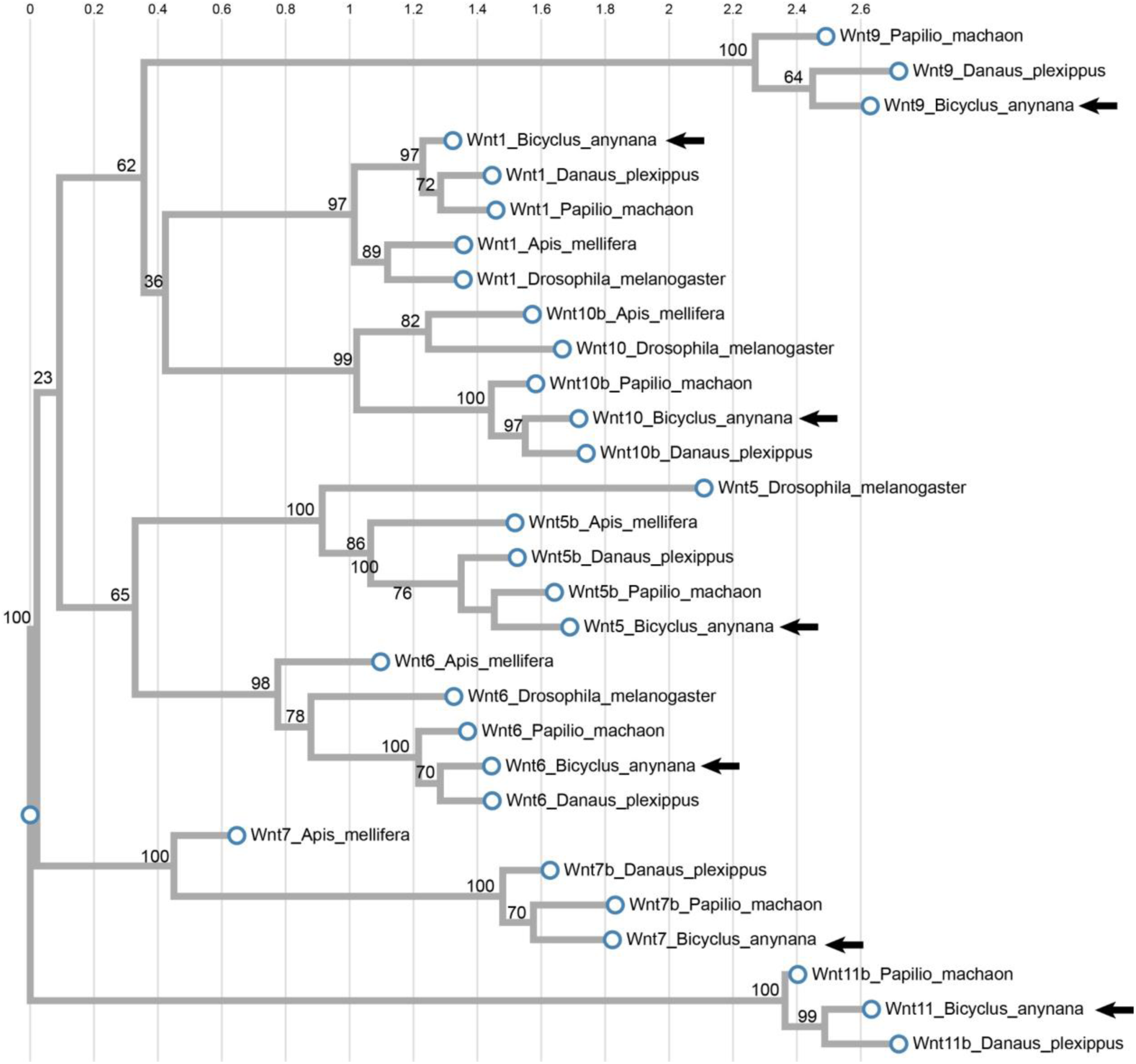
Phylogenetic tree of *wnt* genes created using RAxML. The tree clusters the Wnt family genes together. Vertical lines indicate mean number of nucleotide substitutions per site. Numbers above branches represents bootstrap branch support. Arrows point to the seven *wnt* genes in the genome of *B. anynana*.

**Figure S4.**
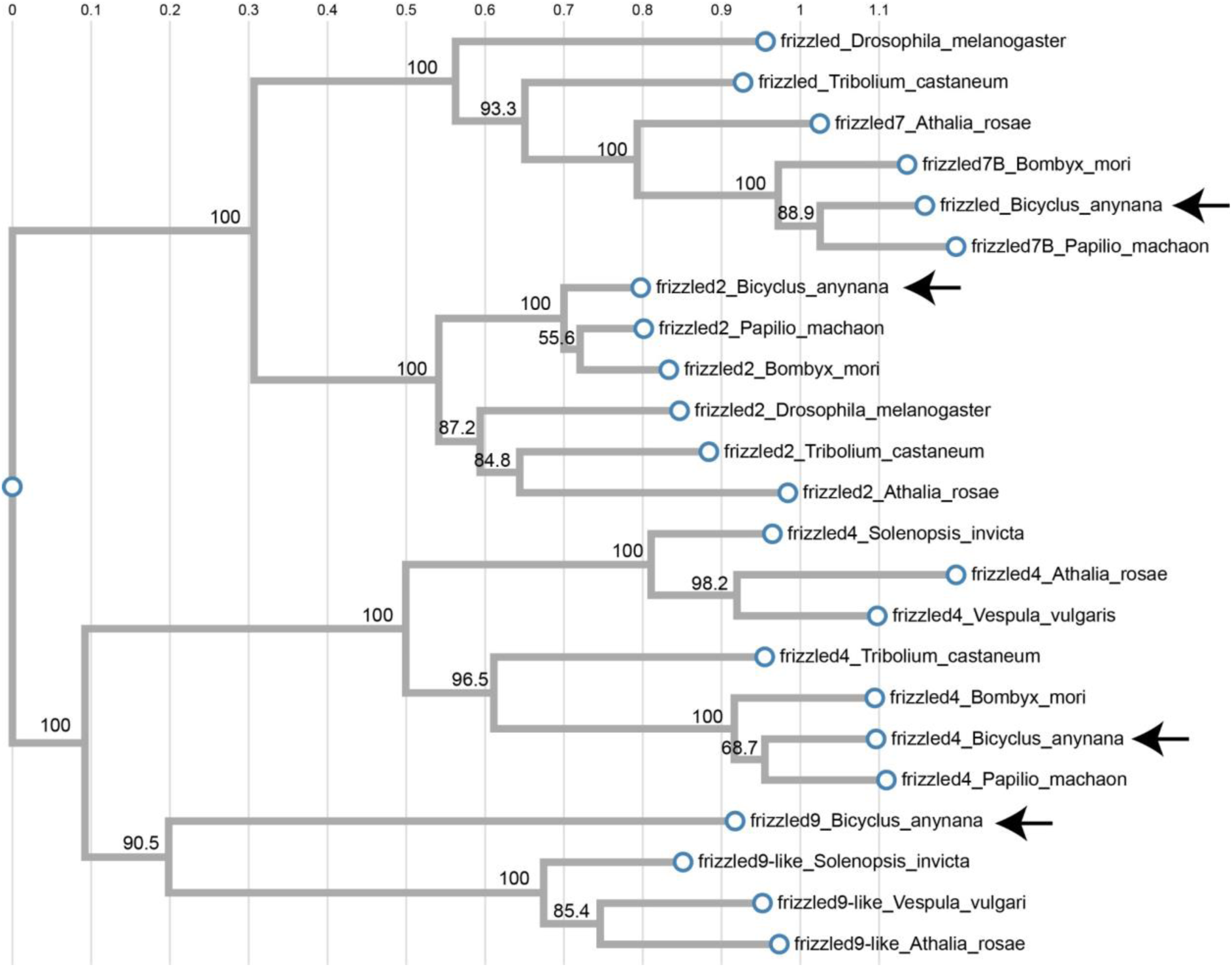
Phylogenetic tree of *frizzled* genes created using fasttree (Maximum likelihood). The tree clusters the frizzled family genes together. Vertical lines indicate mean number of nucleotide substitutions per site. Numbers above branches represents bootstrap branch support. Arrows point to the four *frizzled* genes in the genome of *B. anynana*.

**Figure S5.**
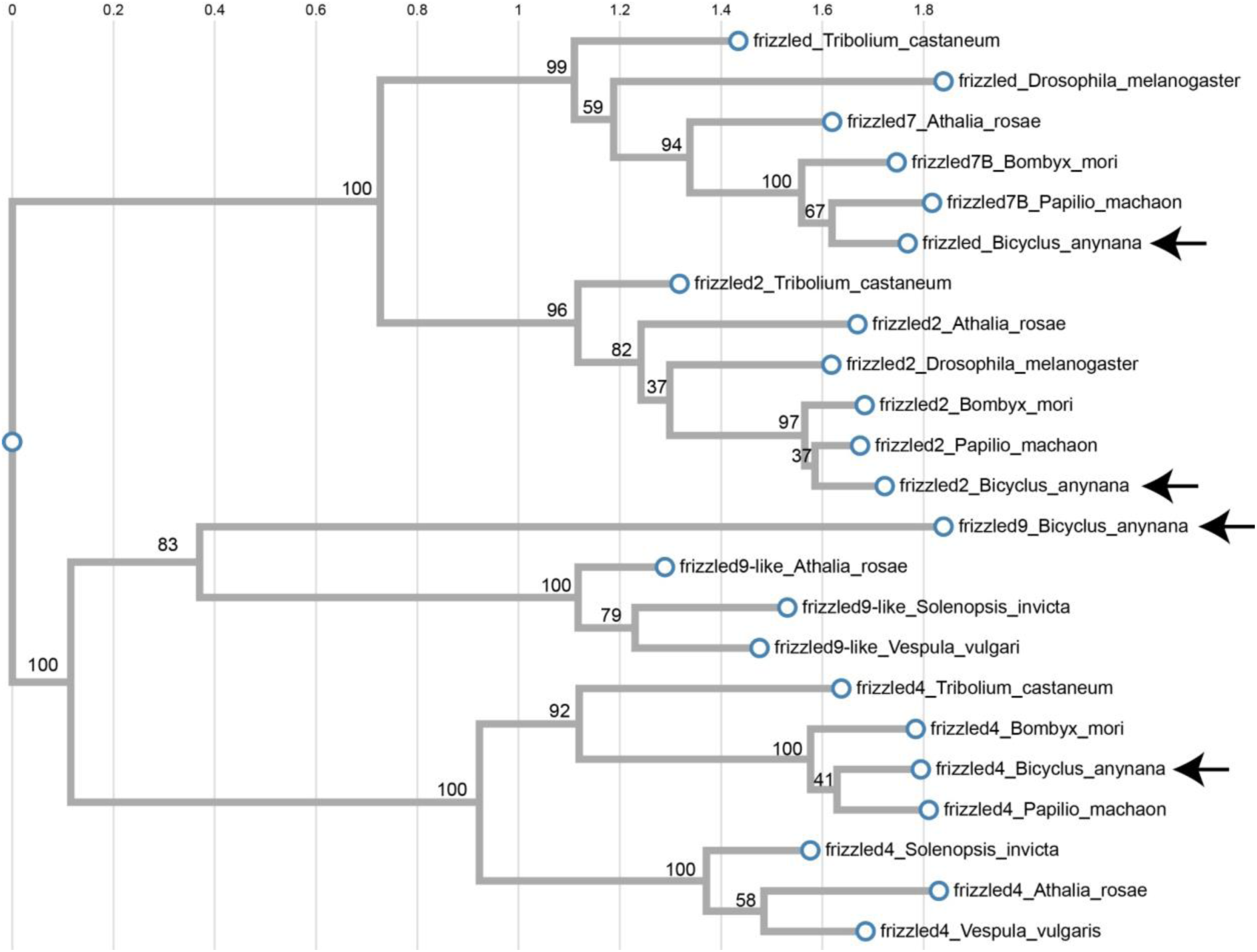
Phylogenetic tree of *frizzled* genes created using RAxML. The tree clusters the frizzled family genes together. Vertical lines indicate mean number of nucleotide substitutions per site. Numbers above branches represents bootstrap branch support. Arrows point to the four frizzled genes in the genome of *B. anynana*.

**Figure S6.**
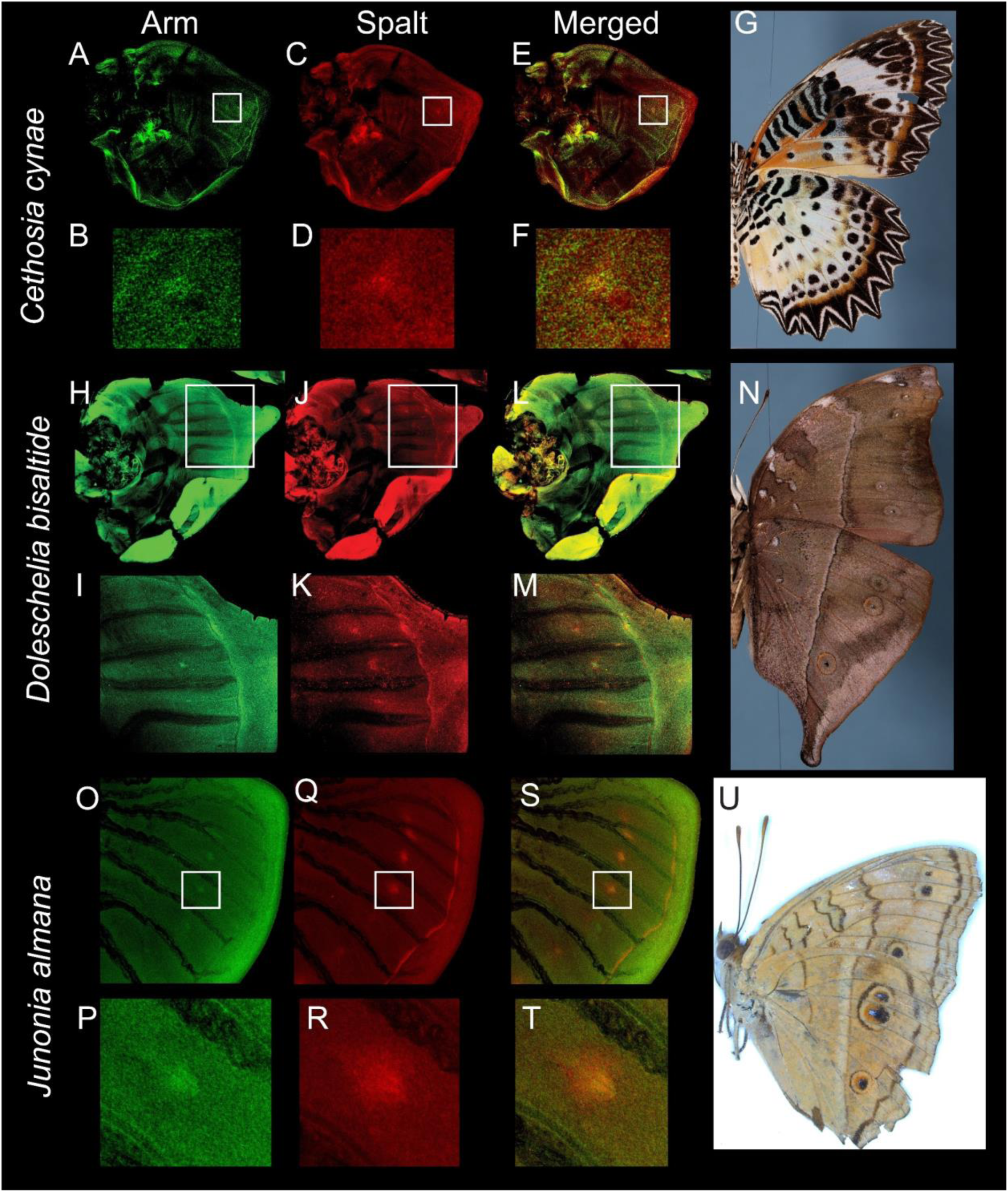
Expression of Arm and eyespot marker gene Spalt in *Cethosia cynae*, *Doleschelia bisaltide* and *Junonia almana* larval wings. Both (**A, B**) Arm and (**C, D**) Spalt are expressed in the central cells of *Cethosia cynae* larval wing. (E and F) Merged expression of Arm and Spalt. (**G**) WT *Cethosia cynae* adult. (**H, I**) Arm and (**J, K**) Spalt are expressed in the central cells of *Cethosia cynae* larval wing. (**L, M**) Merged expression of Arm and Spalt. (**N**) WT *Doleschelia bisaltide* adult. (**O, P**) Arm and (**Q, R**) Spalt are expressed in the central cells of *Junonia almana* larval wing. (**S, T**) Merged expression of Arm and Spalt. (**U**) WT *Junonia almana* adult.

**Figure S7.**
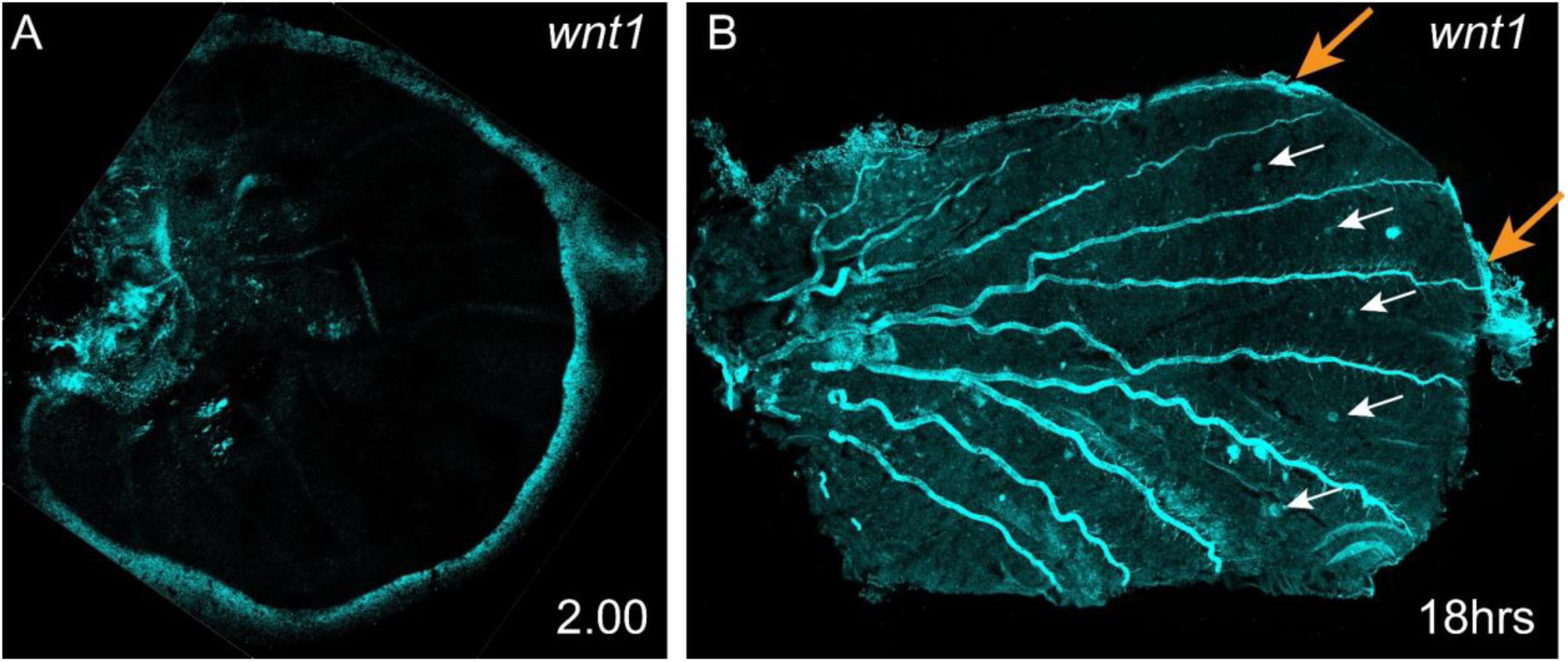
Expression of Arm and *wnt1* in the pupal wing and late larval wing. (**A**) In the late larval wing (2.00) *wnt1* continues to express in the wing margin alone, and not in the eyespot foci. (**B**) In a 18 hrs pupal wing *wnt1* is expressed in the eyespot centers (the foci) and in the wing margin.

**Figure S8.**
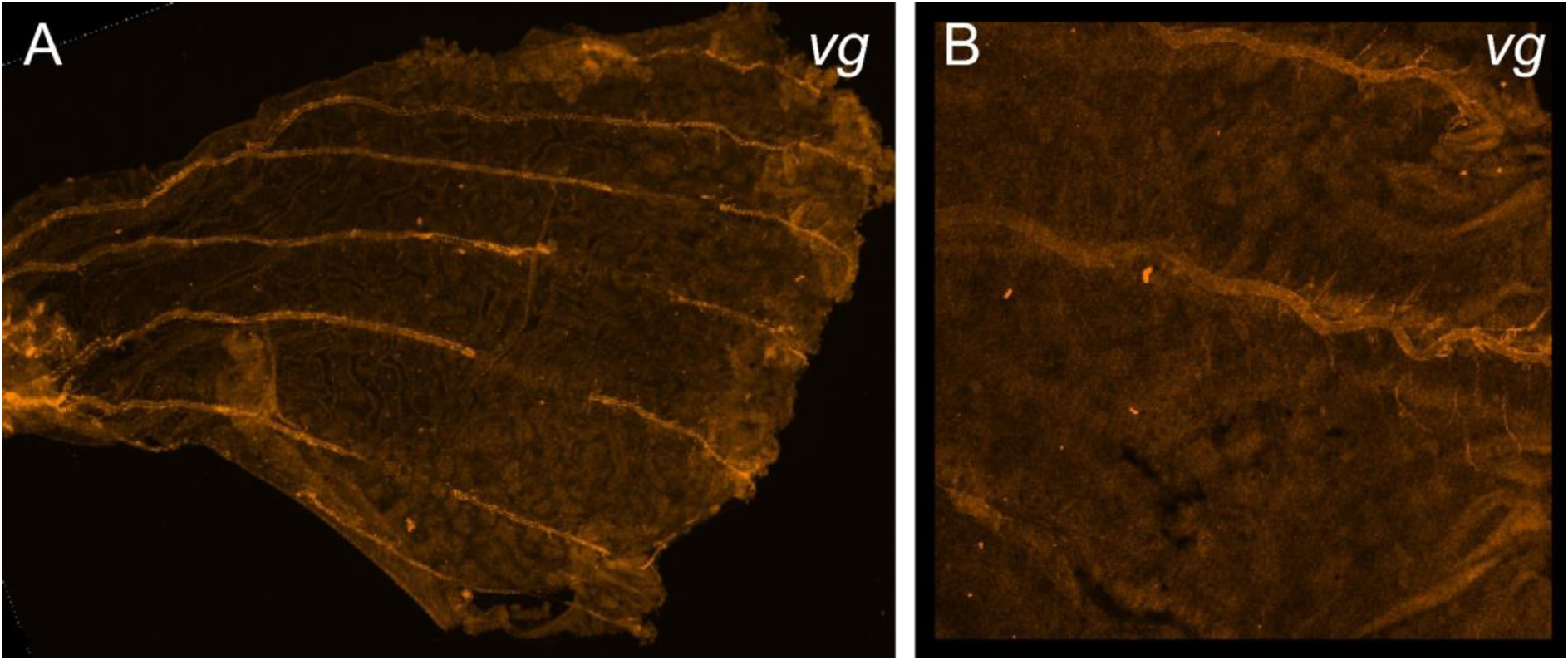
Expression of *vg* in in the pupal wing. No specific domain of expression was observed for *vg* in 18-24 hrs pupal wings.

**Figure S9.**
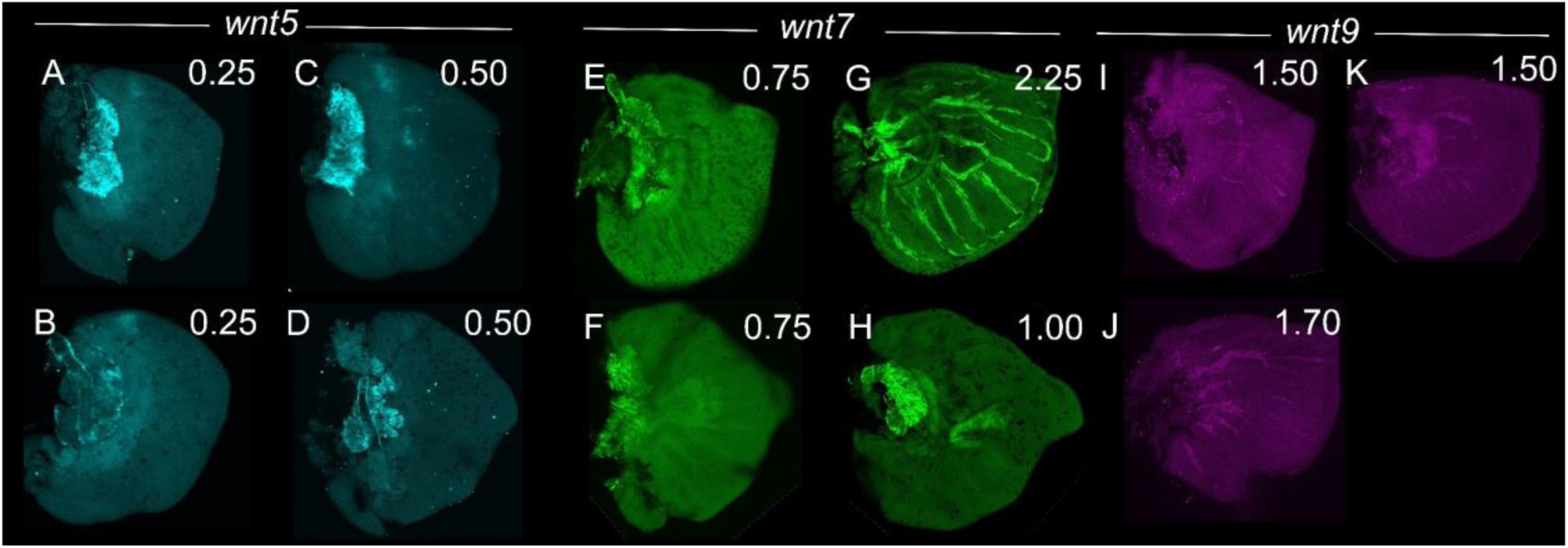
No specific expression of *wnt5, wnt7* and *wnt9* in the larval wings of *Bicyclus anynana*.

**Figure S10.**
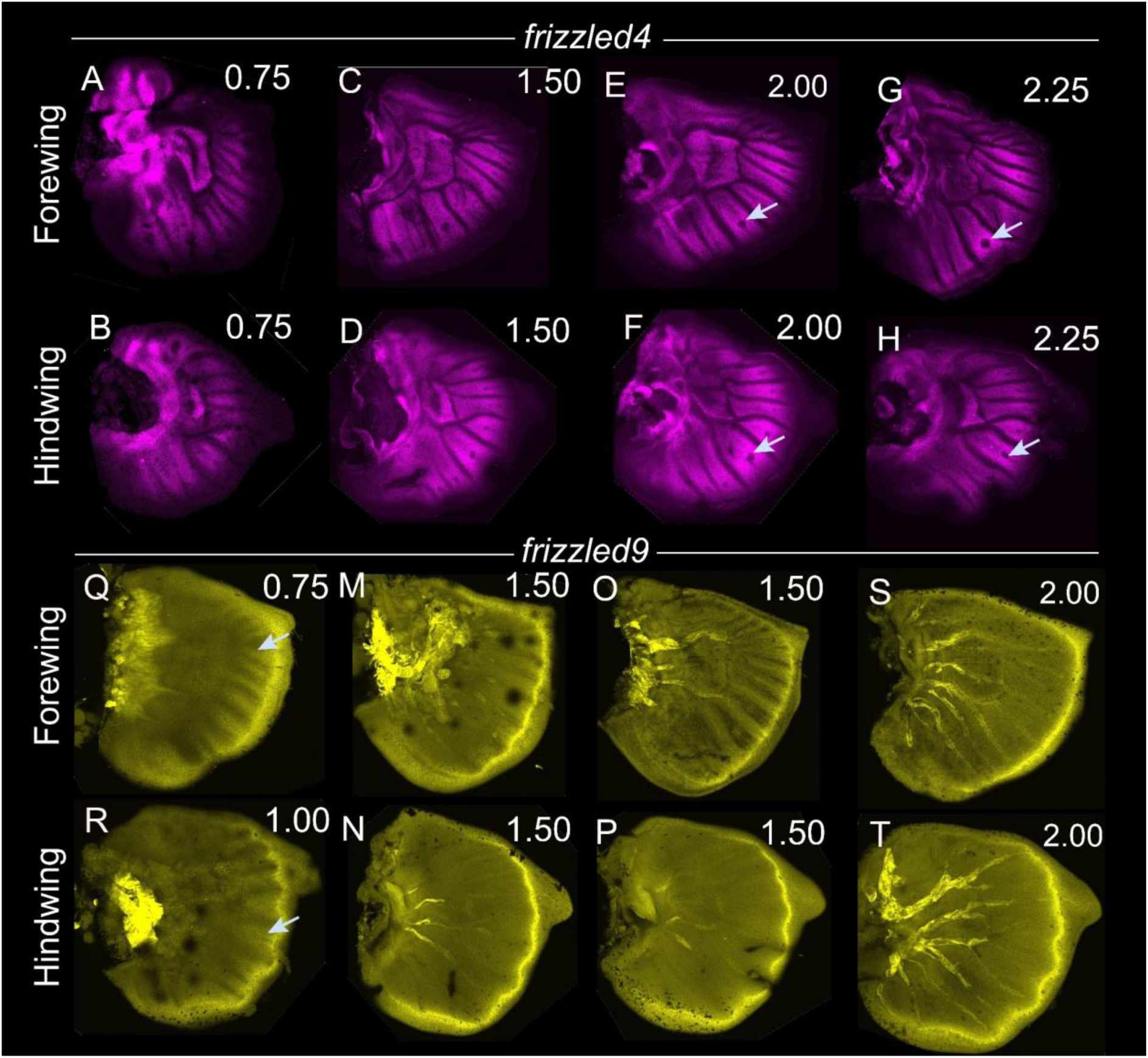
Expression of *frizzled4* and *frizzled9* in the larval wings of *Bicyclus anynana*. (**A-H**) Expression of *frizzled4* in the developing larval wings. During the early larval stage *frizzled4* is expressed in the intervein cells. As the wing develops, *frizzled4* continues to express in the intervein cells but is down-regulated in the eyespot centers. **(Q-T)** Expression of *frizzled9* in larval wings is present initially along the wing margin and in broad fingers projecting from the margin (arrows in Q and R), which become less visible as the wing ages.

**Figure S11.**
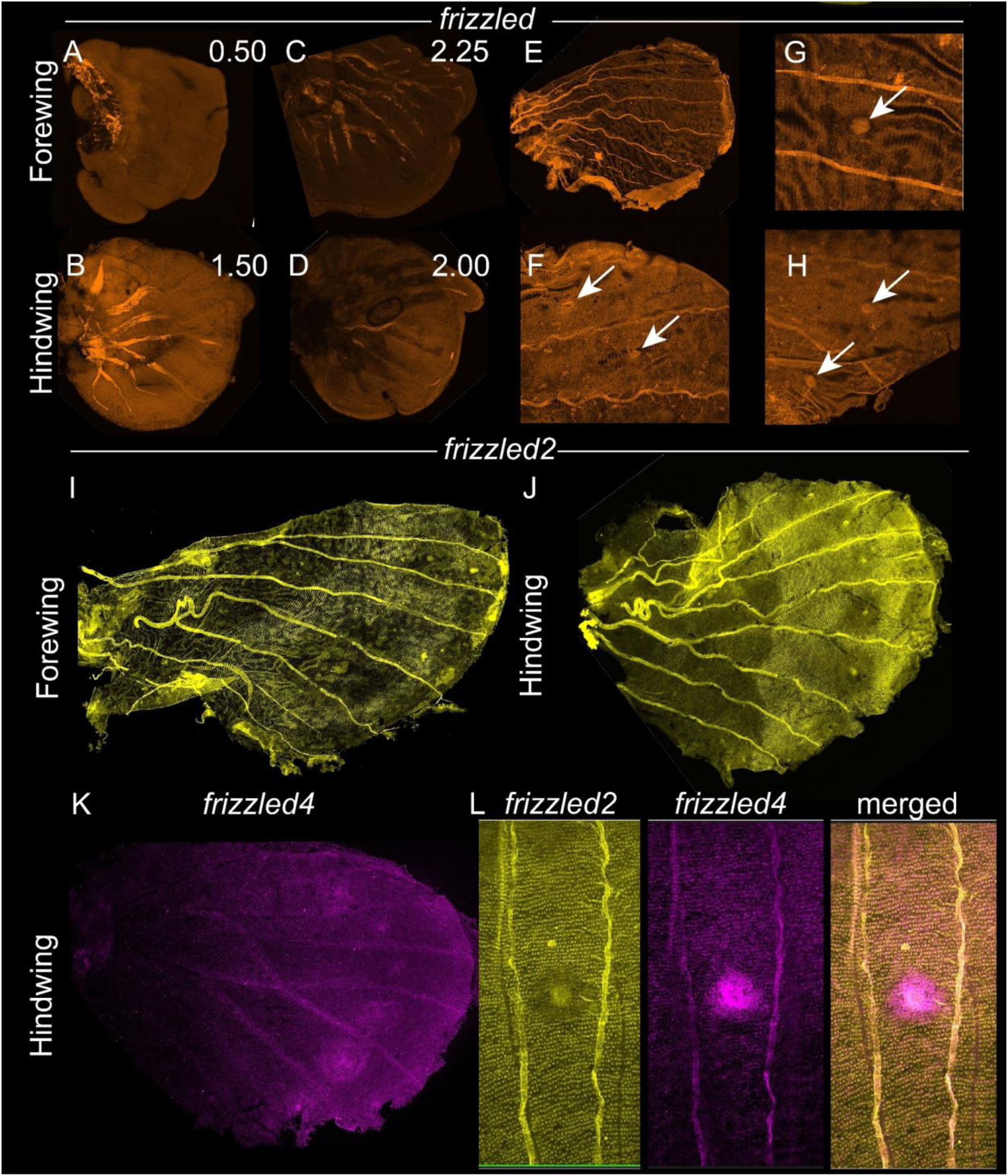
Expression of *frizzled, frizzled2,* and *frizzled4* in the larval and pupal wings of *Bicyclus anynana*. (**A-D**) There is no expression of *frizzled* in the developing larval wings. **(E-H)** During the early larval stage *frizzled* is expressed in the eyespot centers. (**I-J**) Expression of *frizzled2* in 18-24 hrs pupal wings where *frizzled2* has lower expression in the central symmetry system, in the eyespot field, and in the wing margin. (**K**) Expression of *frizzled4* in a 18-24 hr pupal wing showing stronger expression in the eyespot center and in the eyespot field. (**L**) Anti-colocalized expression of *frizzled2* and *frizzled4* in the eyespot field.

**Figure S12.**
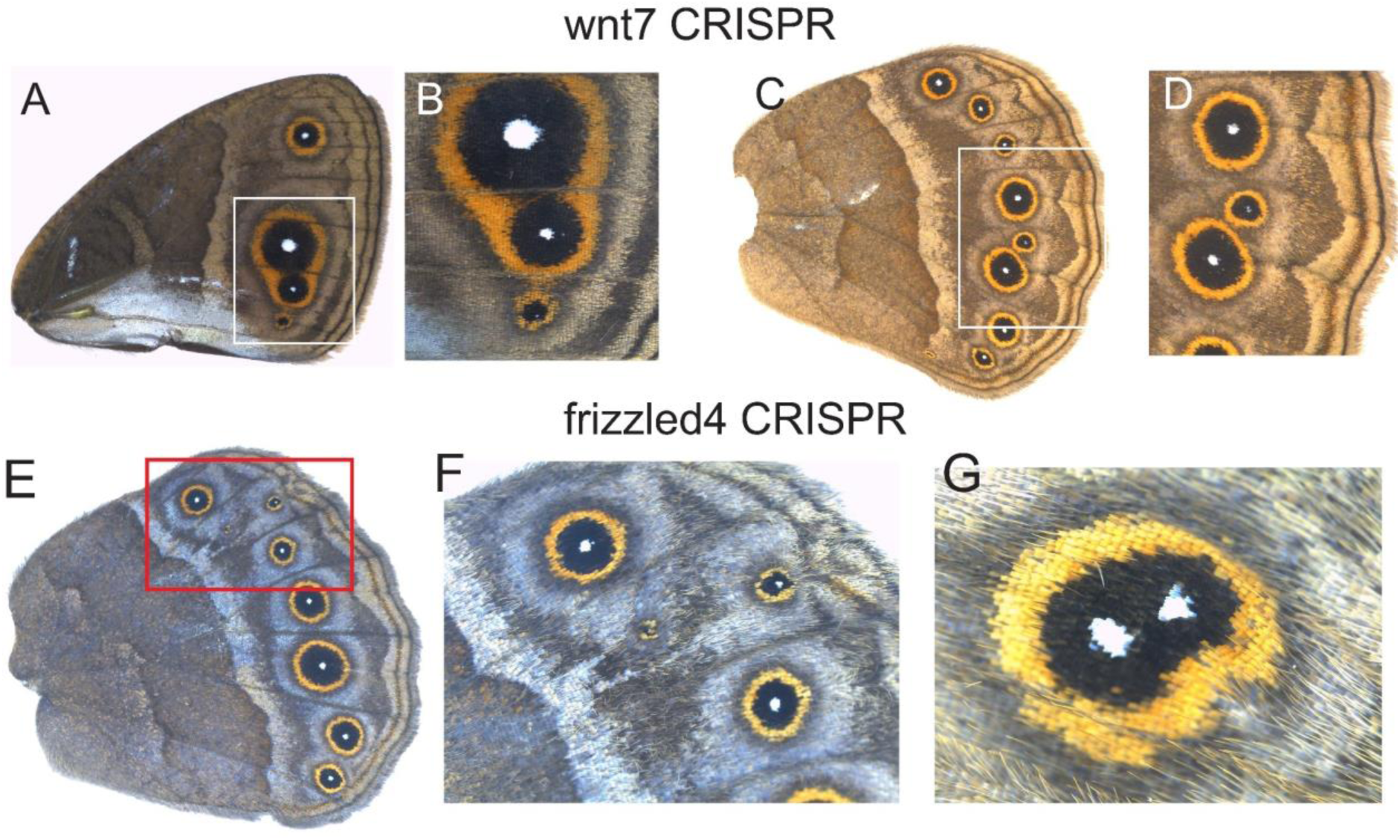
CRISPR-Cas9 on *wnt7* and *frizzled4* in *Bicyclus anynana*. (**A-H**) Knock-out of *wnt7* resulted in venation defects in both forewings and hindwings. These defects resulted in additional eyespots forming in each novel wing sector. (**E-G**) *frizzled4* knockout resulted in two eyespot centers differentiating in the same wing sector.

**Table S1:**
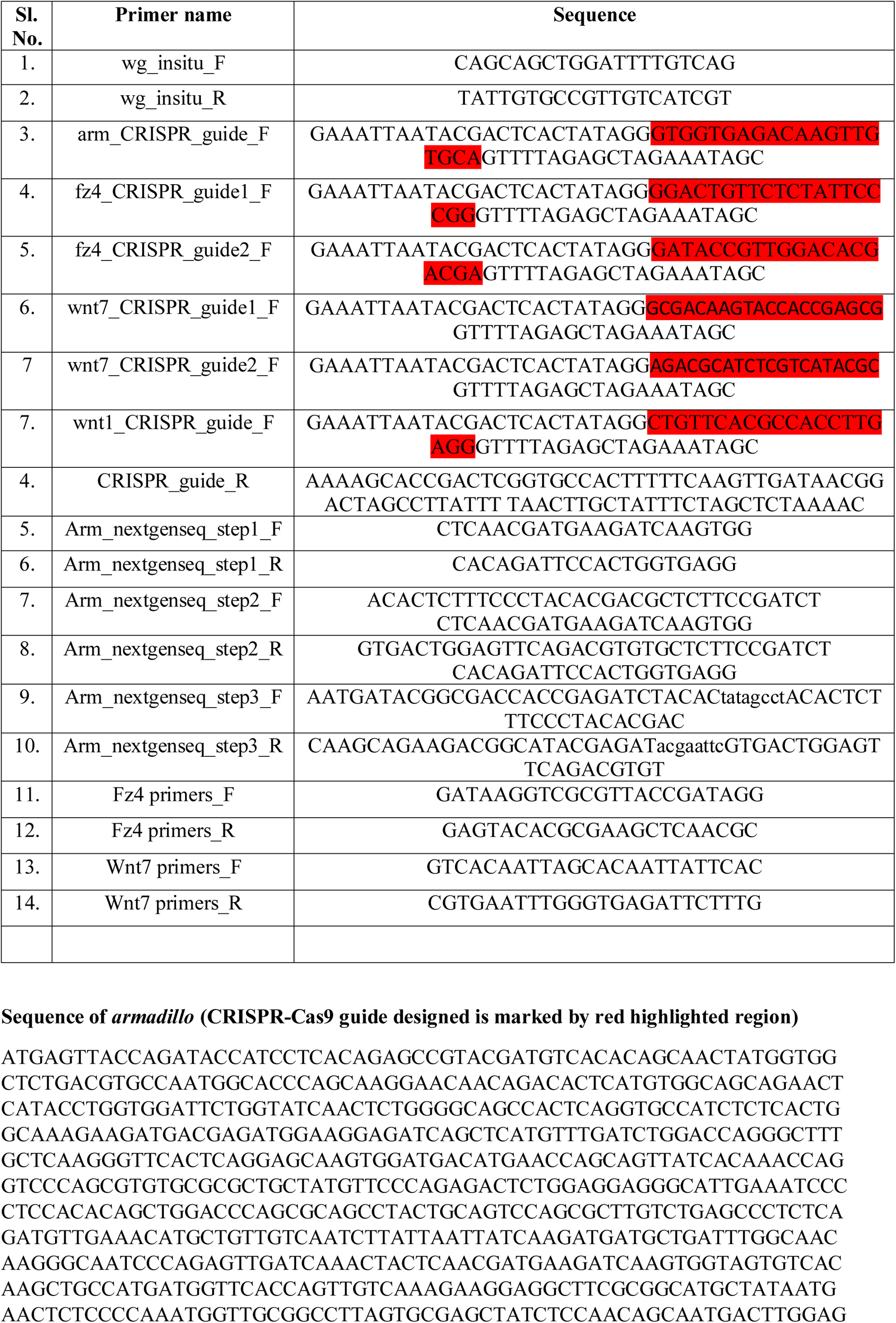
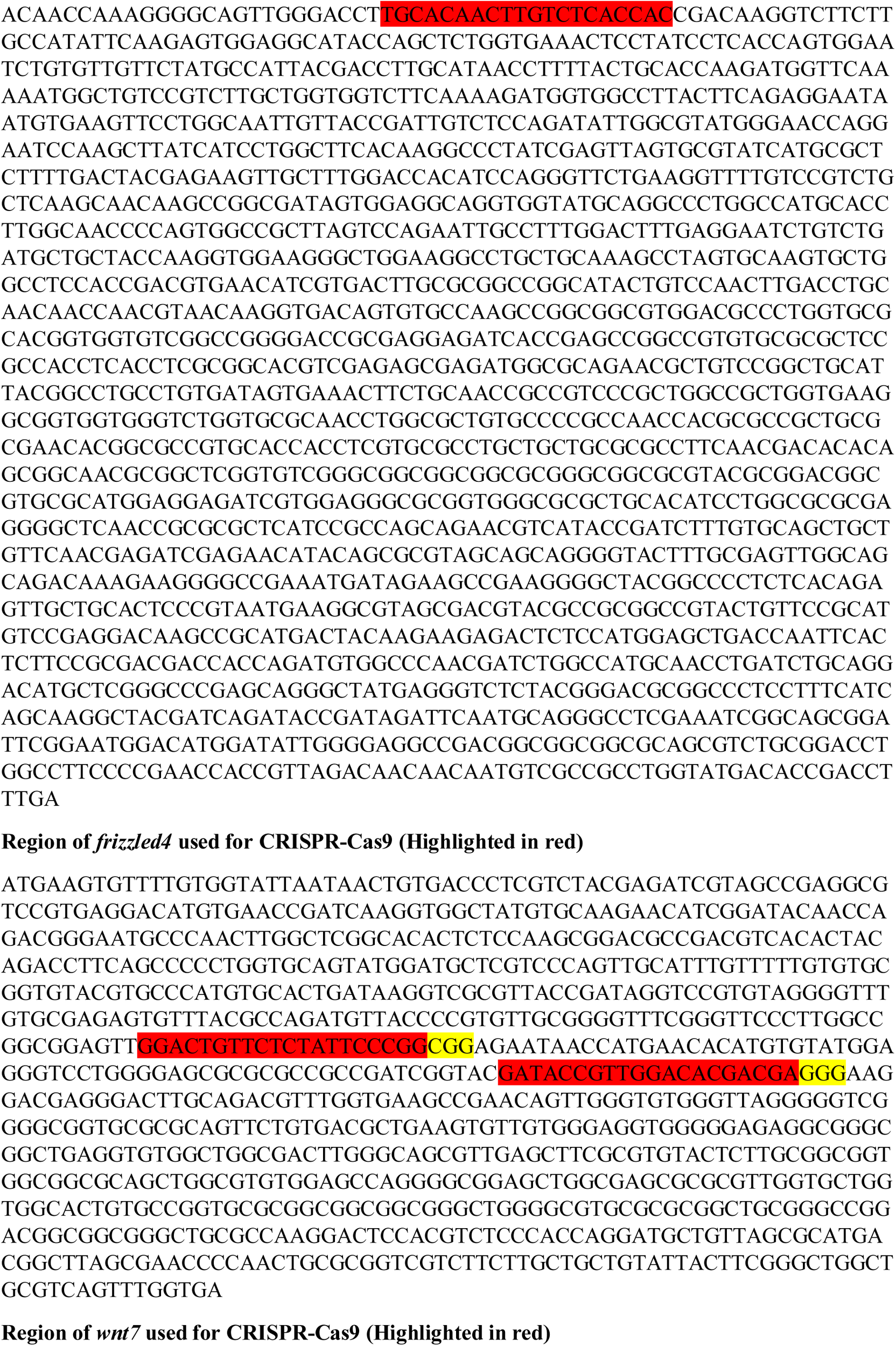
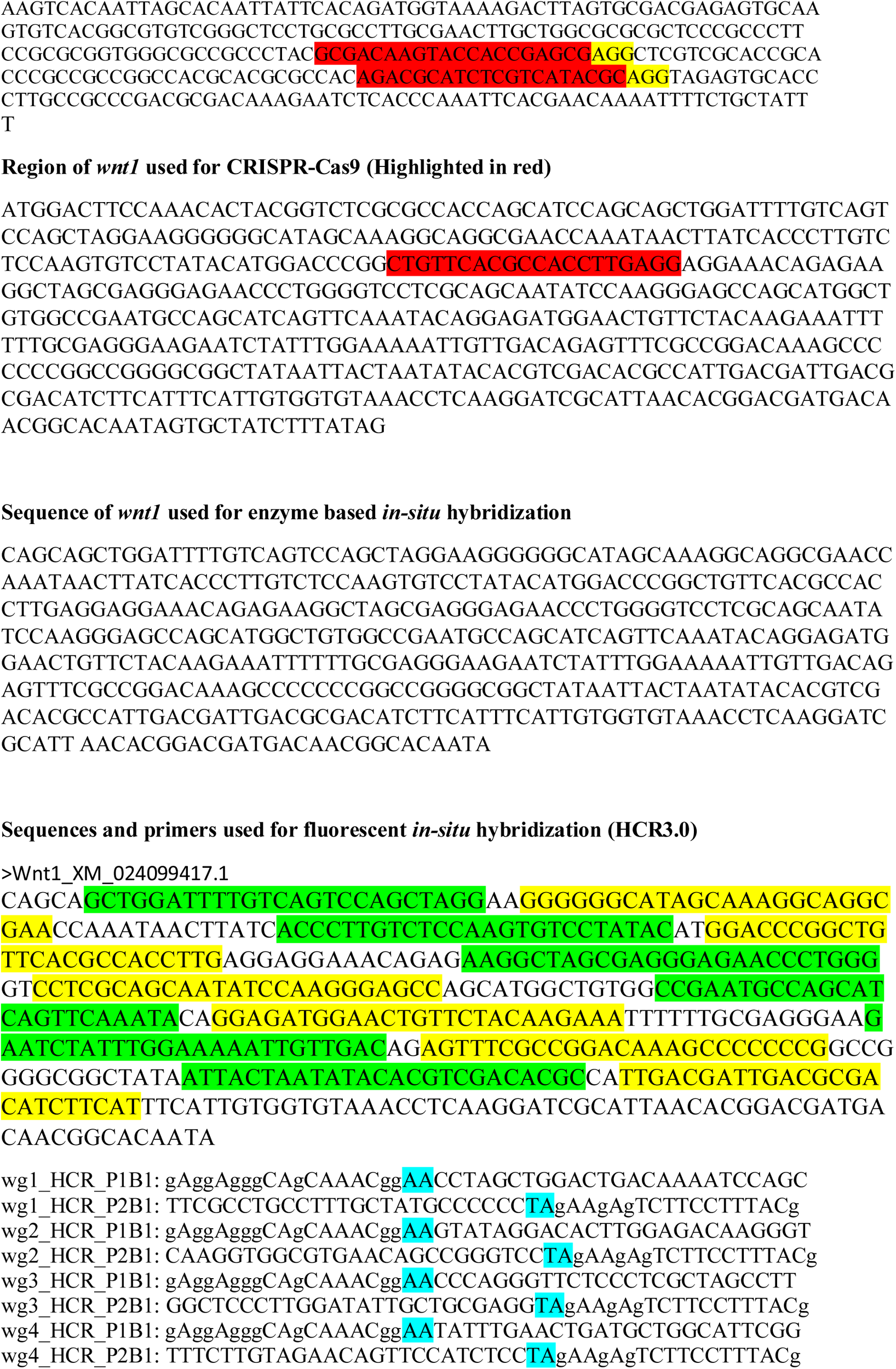
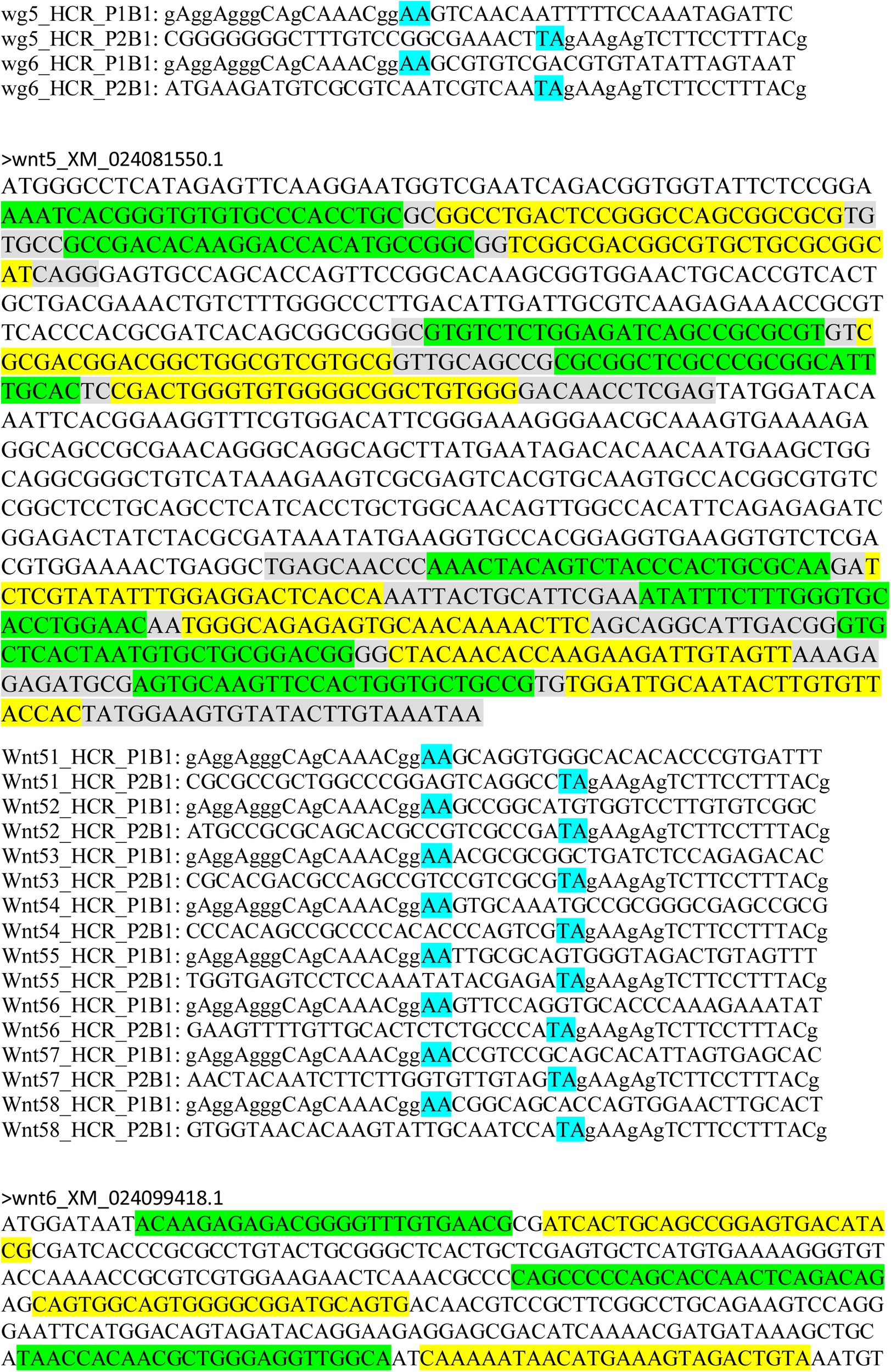
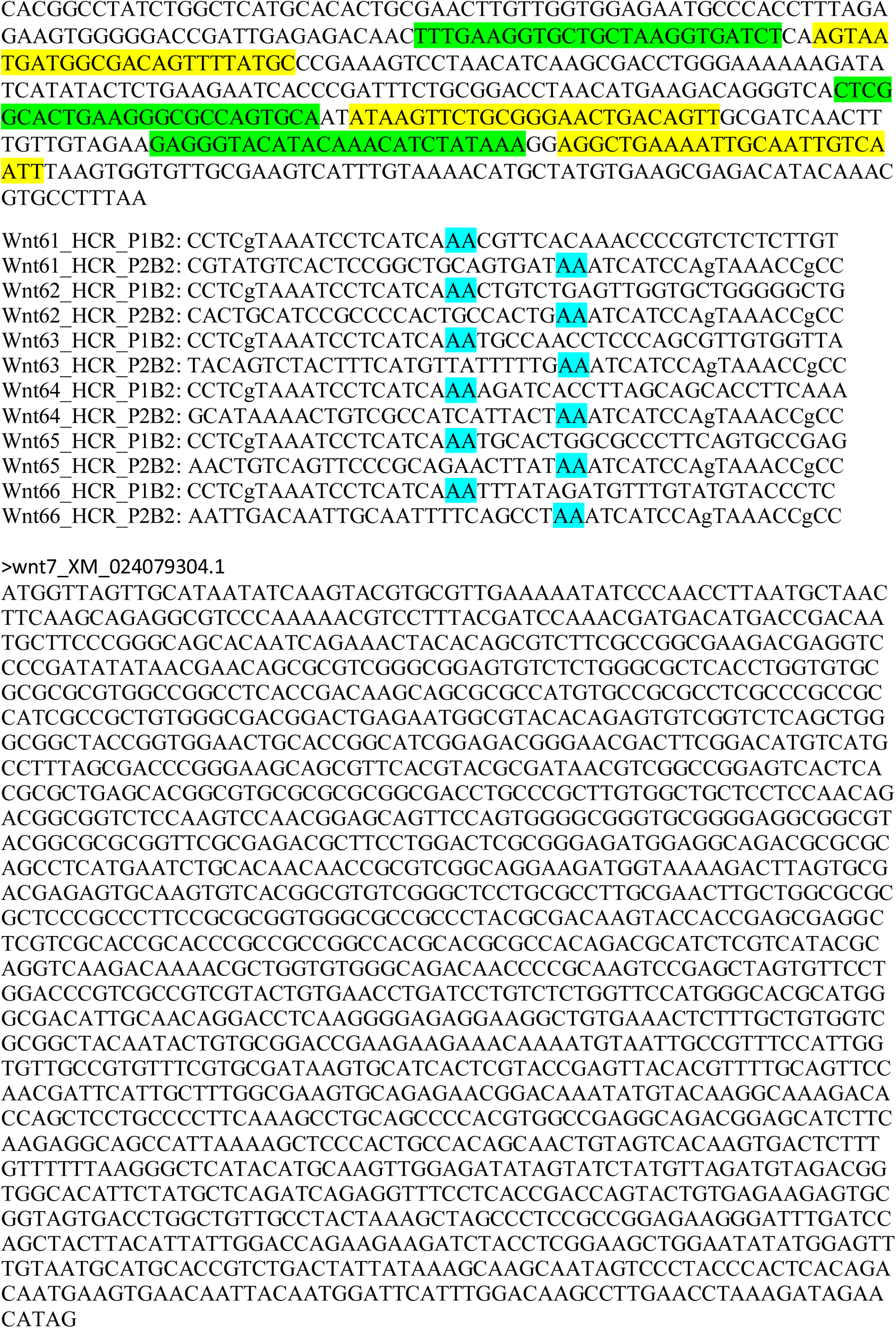
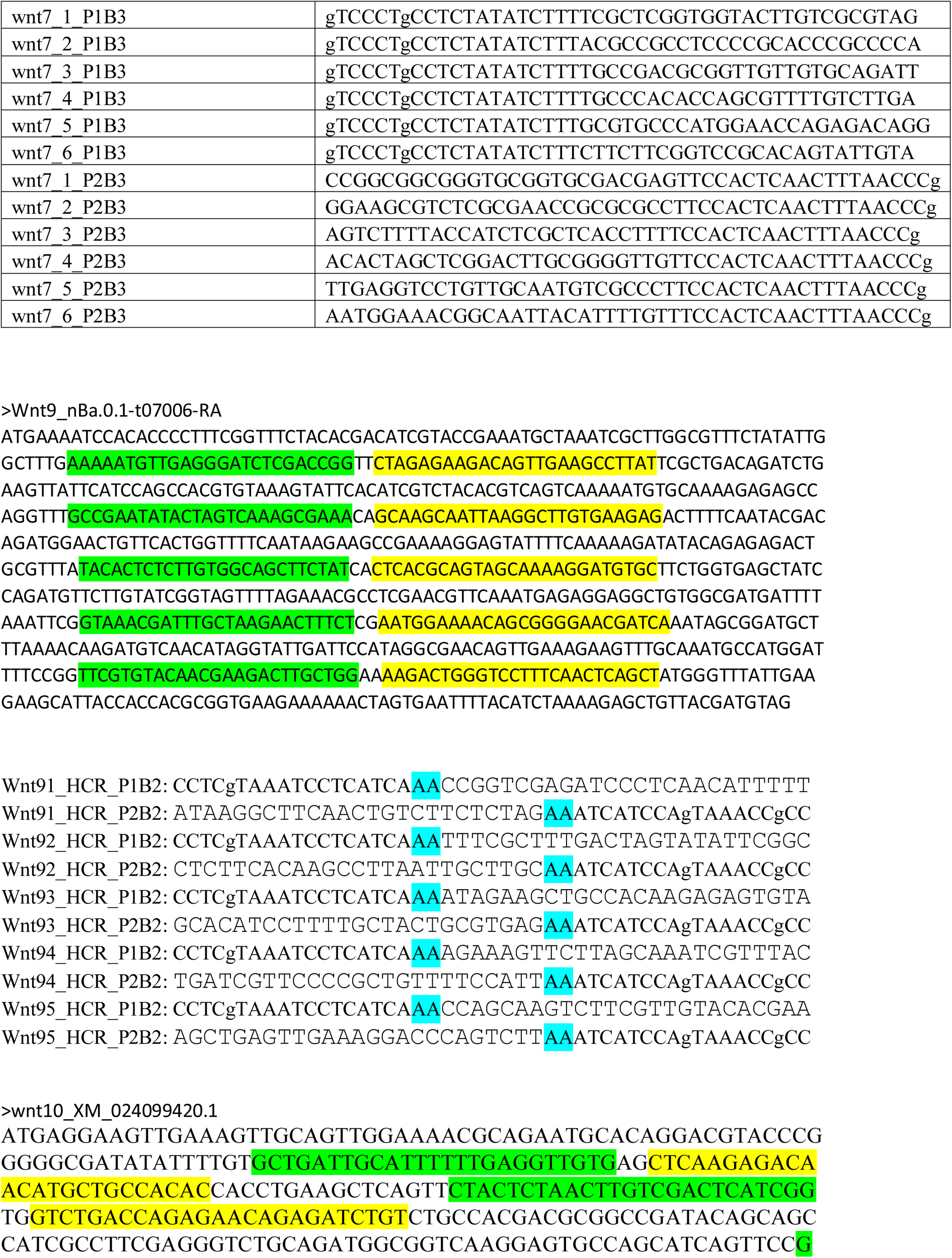
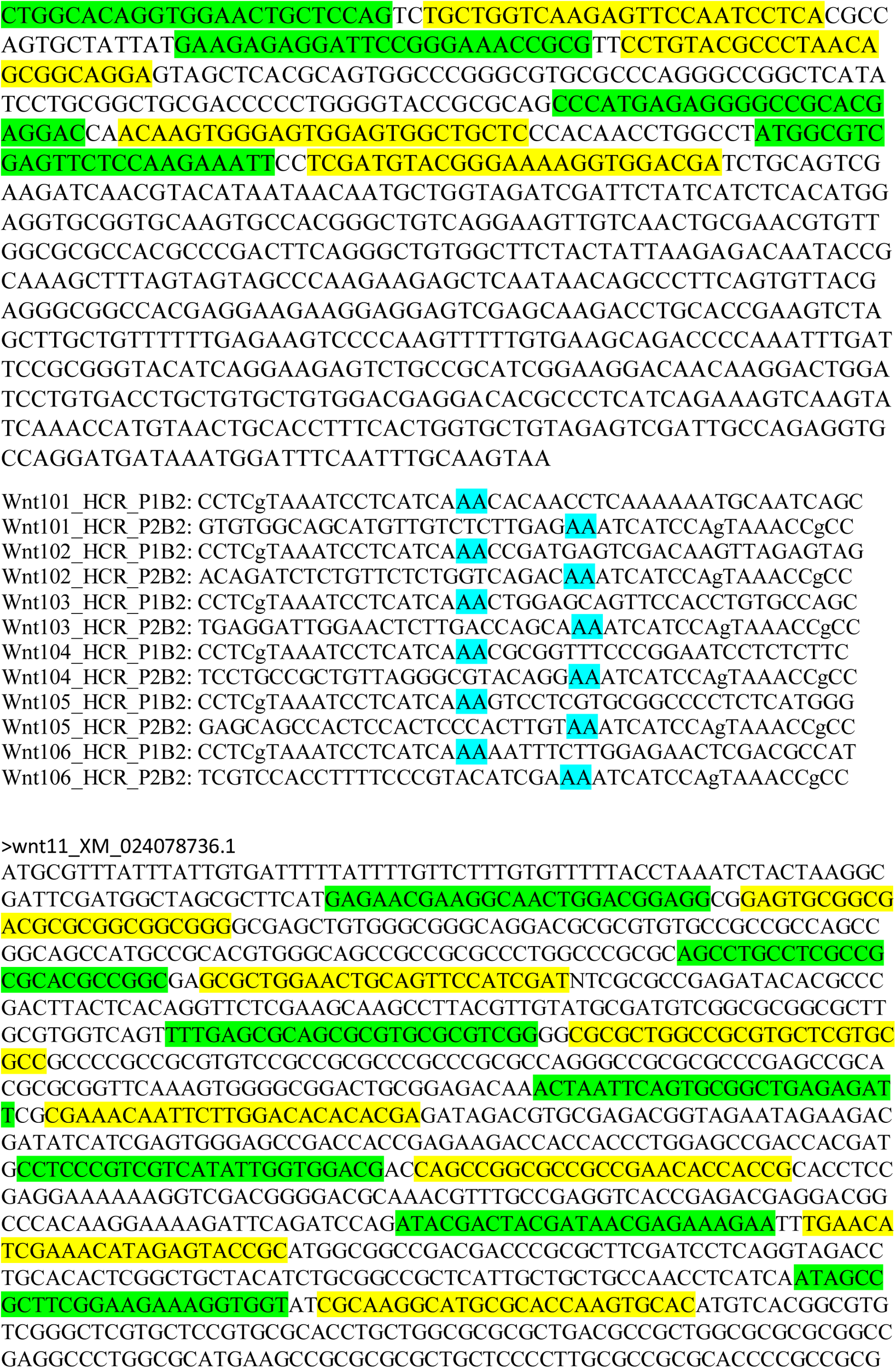
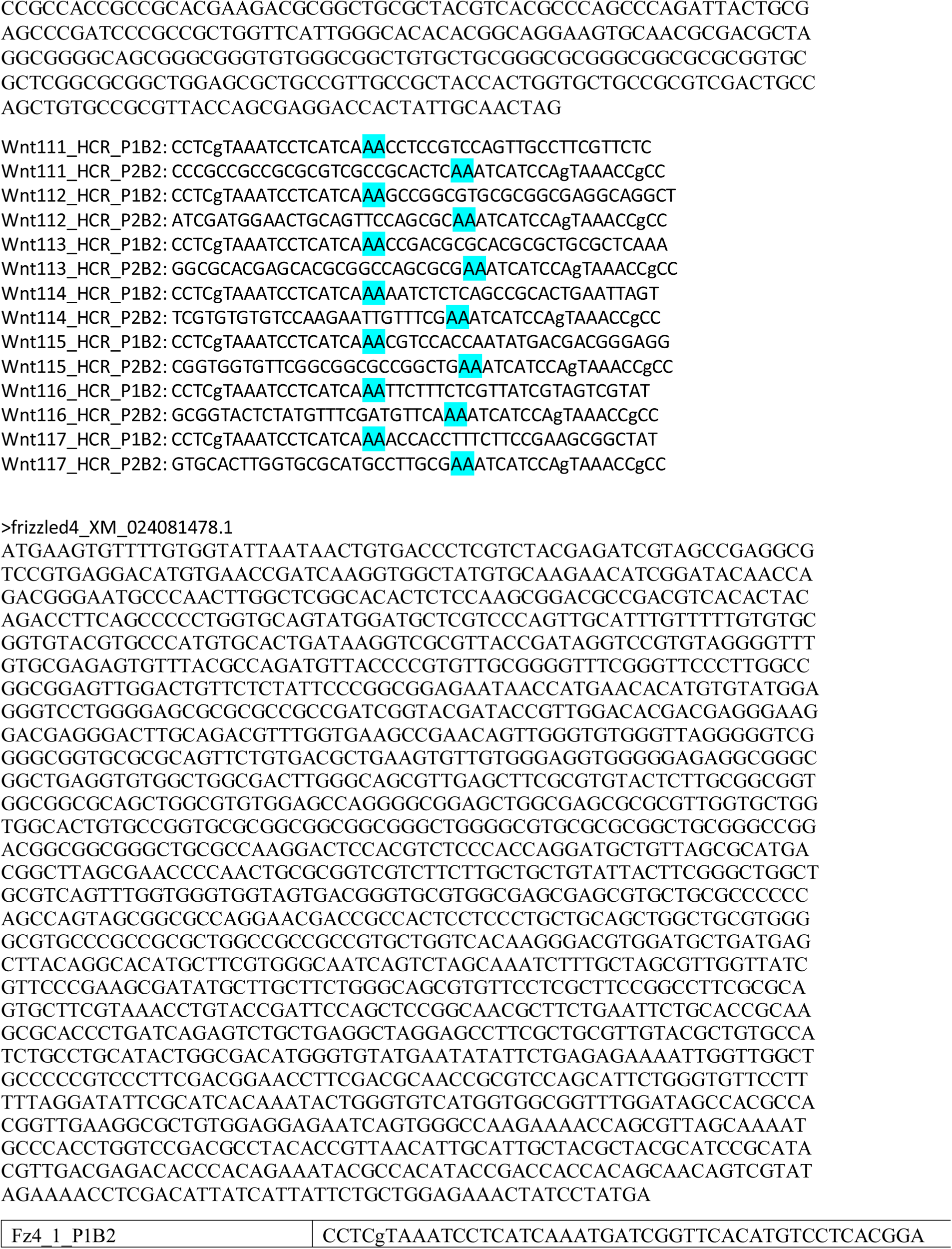
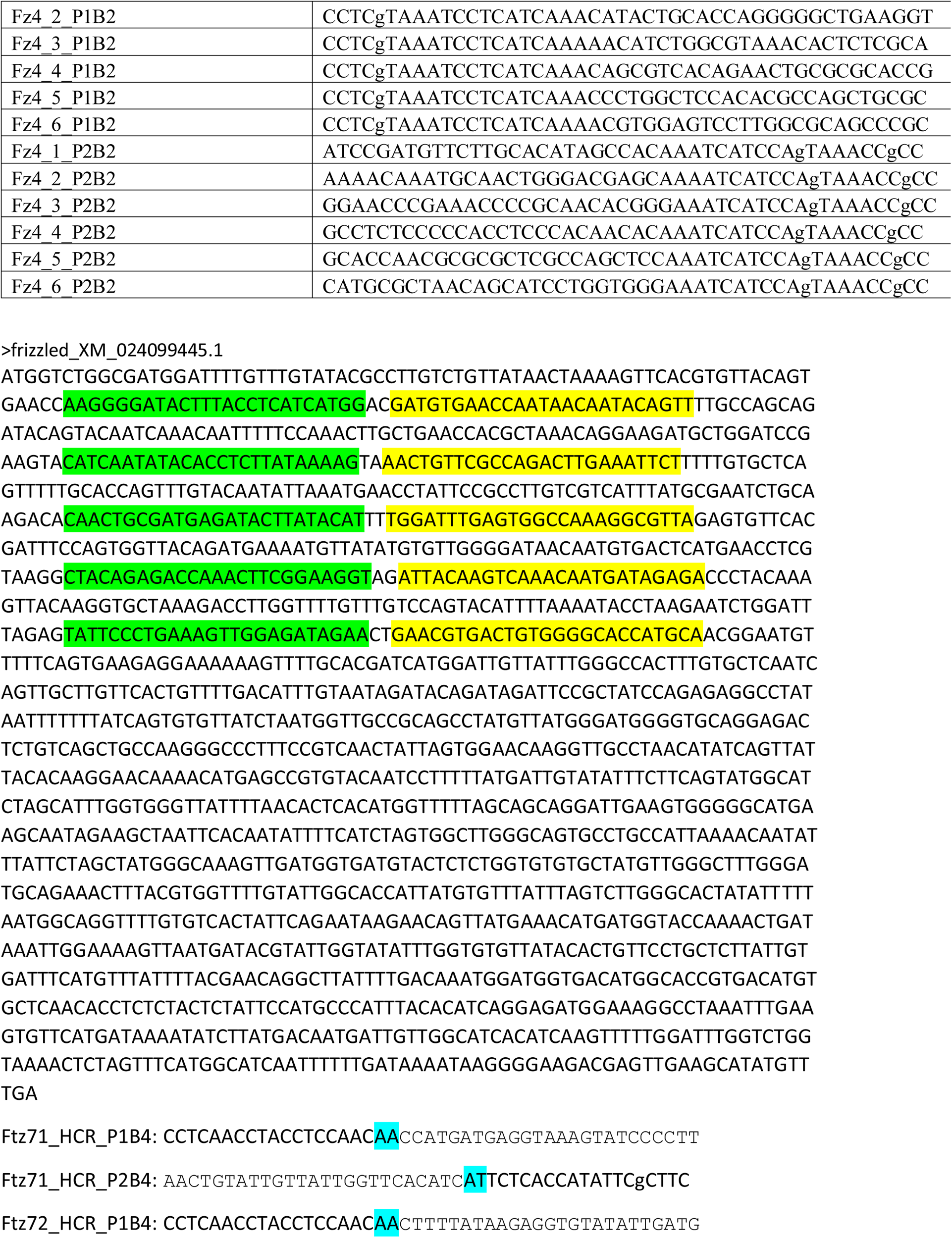
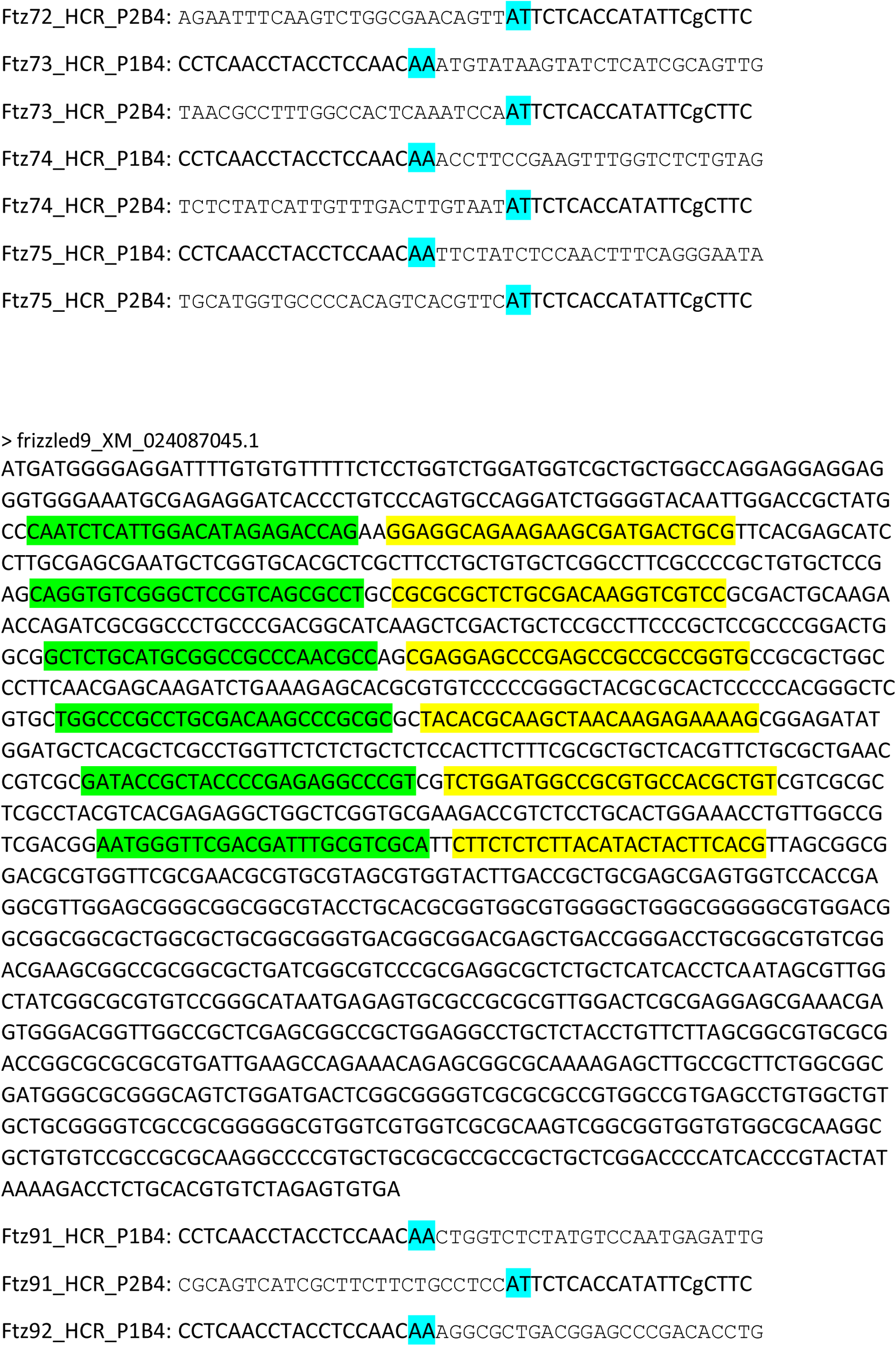
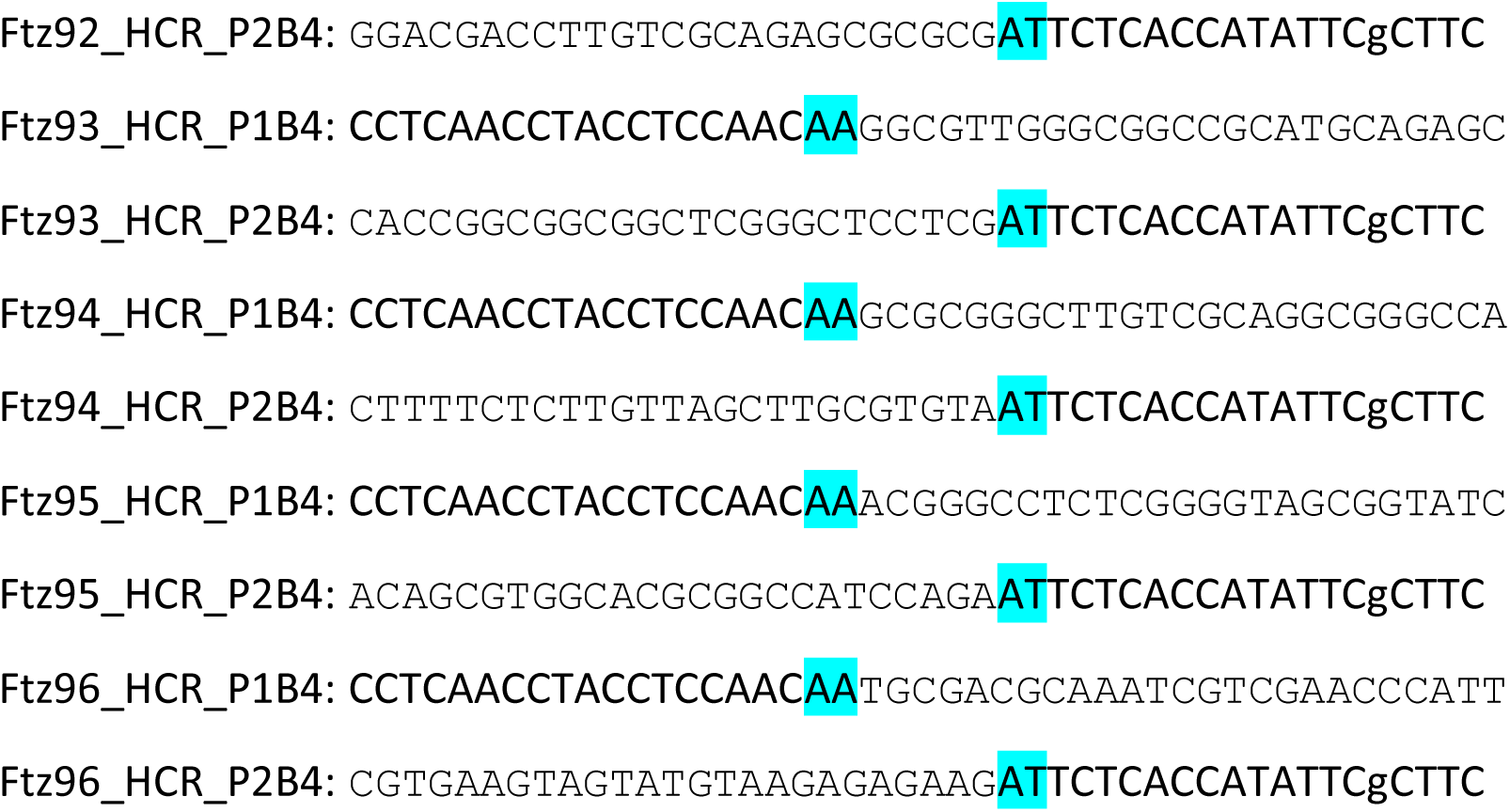
Primer table.

